## Supplementary material for "Transitional evolutionary forms and stratigraphic trends in chasmosaurine ceratopsid dinosaurs : evidence from the Campanian of New Mexico": SUPP TXT1 - Additional information and taxonomic review

### Supplementary figure captions

#### Figure S1

**Hypothesis of chasmosaurine relationships from Lehman (1998).** Lehman (1998) proposed that Campanian chasmosaurines belonged to two lineages evolving divergently from initially similar morphology. A *Chasmosaurus* lineage (left, *C. russelli* **1-3**, *C. belli* **4-7**) was characterized by progressive shallowing of the parietal median embayment, development of epiparietals into an elongate ridge at locus ep1, and lateral migration of loci ep2 and 3. In contrast, an *Agujaceratops* (**8**) - *Pentaceratops* (**9**, **10**) - *Anchiceratops* (**11**, **12**) lineage (right) was characterized by a deepening median embayment which caused rotation of epiparietals at locus ep1 to form the butterfly-wing orientation characteristic of *Anchiceratops*. Lehman's hypothesis was consistent with the stratigraphic distribution of the depicted specimens, and was further supported by the subsequent discovery of new taxa. Facsimile of Lehman (1998).

#### Figure S2

**Alternative hypotheses of chasmosaurine phylogeny.** Two recent phylogenetic analyses contrast the dual-lineage hypothesis of Lehman (1998). **A**, Sampson et al. (2010) recover a phylogeny where *Pentaceratops* is unrelated to *Anchiceratops*; where *Vagaceratops* is unrelated to *Chasmosaurus*; and where instead *Kosmoceratops* and *Vagaceratops* form an outgroup to *Anchiceratops* and all other more derived chasmosaurines. **B**, Mallon et al. (2014; drawing on the same phylogenetic matrix as Sampson et al., 2010) published a phylogeny where the Lower Maastrichtian taxa *Anchiceratops* and *Arrhinoceratops* occur in a basal polytomy, and some of the stratigraphically oldest taxa form the most derived clade (Middle to Upper Campanian *C. belli* + *C. russelli*). These new analyses require significant ghost lineages be present for most clades, for which there is currently no fossil evidence. **A**, **B** adapted from Sampson et al. (2010) and Mallon et al. (2014) respectively.

#### Figure S3

**Historical stratigraphic terminology of the Fruitland and Kirtland Formations.** The Fruitland and Kirtland Formations of the San Juan Basin have undergone many changes regarding terminology, and (more importantly) the definitions of member or formational contacts. Here I illustrate all the revisions in chronological order, ending with the current terminology and definitions used in this study. Adapted from Bauer (1916); Baltz et al. (1966); Fassett and Hinds (1971); Hunt and Lucas (1992; 2003); Sullivan et al. (2005) and Lucas et al. (2006). Section thickness in meters (m).

#### Figure S4

**Taxon A holotype SMP VP-1500 parietal exposed before collection and preparation**. (**A**) An area of rugose concreted bone occurred at locus **ep1** and extended around the anterior border of the median embayment (**em**), but was mostly unrecoverable during preparation. This is interpreted here representing ep1. (**B**) The median bar (**mb**) of SMP VP-1500 was slightly displaced laterally and ventrally. **ep1**, epiparietal 1. **f**, parietal fenestra. **L-lr / R-lr**, Left / Right lateral rami of the posterior bar. Hammer for scale = 28 cm.

#### Figure S5

**Taxon A holotype SMP VP-1500 isolated frill epiossification**. Isolated frill epiossification found with parietal. Possibly an epiparietal, although squamosal fragments were found weathered out on the surface. Scalebar = 5 cm.

#### Figure S6

**Taxon A holotype SMP VP-1500 squamosal fragments.** Two of the largest fragments of the squamosal found weathered on the surface next to the parietal. **A** (dorsal)**, B** (ventral; inferred orientations), squamosal fragment exhibiting two fused episquamosals (**es**). **C, D**, elongate fragment, possibly from the posterior end of the squamosal where it narrows. Both fragments exhibit characteristic ceratopsian vascular surface texture. Scalebar = 10 cm.

#### Figure S7

**Taxon A holotype SMP VP-1500 jugal.** Possible ventralmost end of a jugal with fused epijugal (**ej**), in lateral and medial views (inferred). Alternatively, this may be an episquamosal fused to a small part of the squamosal. Scalebar = 5 cm.

#### Figure S8

**Taxon B holotype NMMNH P-27468 right squamosal.** Ventral (left) and dorsal (right) views. **sb**, squamosal bar. **es**, episquamosal. Scalebar = 10 cm.

#### Figure S9

**Taxon B holotype NMMNH P-27468 left jugal.** Ventral two thirds of left jugal with fused epijugal (**ej**) and partial quadratojugal in posterior (**A**), left lateral (**B**) and Anterior (**C**) views. A size comparison of NMMNH P-27468 (**D**) is compared with *Utahceratops gettyi* referred specimen UMNH VP-12198 (**E**). Scalebars = 10 cm.

#### Figure S10

**Angle formed by the lateral rami of the parietal posterior bar in *Pentaceratops* and close relatives.**

In most chasmosaurines the lateral rami of the parietal posterior bar meet medially at an angle, forming an embayment. Specimens are shown here in stratigraphic order where possible. (**A**) is probably the stratigraphically oldest specimen illustrated (see supp. info. text). The stratigraphically separated taxa *Utahceratops* (**B**), *Pentaceratops* (**C**, **D**, **F**), Taxon A (**G**), to Taxon B (**H**) form a morphologic spectrum, recording overall decrease in the angle of the lateral rami of the posterior bar, deepening and narrowing the median embayment. *Agujaceratops* specimen UTEP P.37.7.065 SDMNH 43470 (**E**) is of uncertain stratigraphic position, but may be roughly equivalent to the Hunter Wash Member of the Kirtland Formation, New Mexico, from which Taxon A (**G**) and Taxon B (**H**) were collected. Specimens not shown to scale (see main text figure for relative sizes).

#### Figure S11

**Angle formed by the lateral rami of the parietal posterior bar in *Chasmosaurus* and close relatives.**

In most chasmosaurines the lateral rami of the parietal posterior bar meet medially at an angle, forming an embayment. Taxa are illustrated in stratigraphic order where possible. The stratigraphically separated taxa "*Chasmosaurus russelli"* (**B-D**), *C. belli* (**E-J**), *Vagaceratops irvinensis* (**K**), and *Kosmoceratops richardsoni* (**H**) form a morphologic spectrum, recording overall increase in the angle of the lateral rami of the posterior bar, shallowing the median embayment. Specimens not shown to scale.

#### Figure S12

**Selected measurements of Taxon A holotype SMP VP-1500**

Parietal shown in dorsal view. Measurements in cm (1.d.p).

#### Figure S13

**Selected measurements of Taxon B holotype NMMNH P-27468**

Parietal shown in dorsal view. Measurements in cm (1.d.p).

#### Figure S14

**Selected measurements of Taxon B holotype NMMNH P-27468**

Jugal - epijugal shown in posterior (left) and left lateral (right) views. Measurements in cm (1.d.p).

#### Figure S15

**Previously undescribed specimen, aff. *Pentaceratops* n.sp., NMMNH P-37880**.

NMMNH P-37880, a partial right lateral ramus of parietal posterior bar in posterior, dorsal, medial, and ventral views. Although an isolated skull fragment, the posterior bar of the parietal is the most diagnostic element in Campanian chasmosaurines. Specimen recovered from the Fossil Forest Member, Fruitland Formation (San Juan Basin, New Mexico) and is morphologically most similar to other specimens referred to aff *P.* n. sp.. Abbreviations: **em**, median embayment of the posterior bar; **ep**, epiparietal loci numbered by hypothesized position (no epiossifications are fused to this specimen). **f**, parietal fenestra. Scalebar equals 10 cm. Reconstruction line drawing based on c.f. *P. sternbergii* specimen UKVP 16100.

#### Figure S16

**Reidentification of parietal median bar of *Bravoceratops polyphemus* (Wick and Lehman, 2013)**

The parietal fragment of TMM 46015-1 (*Bravoceratops polyphemus*, **A**) is reidentified here as representing the anterior half of the parietal median bar as it compares favorably with the anterior median bars of c.f. *Pentaceratops* (USNM 8604, **B**; PMU 24924, **C**) and *Chasmosaurus belli* (CMN 491; **D**). The specimen was previously identified as the posterior half of the parietal median bar by Wick and Lehman (2013, **E**), and interpreted as such bore most of the morphological features which distinguished this new taxon. Scalebars equal 10 cm. No scale available for PMU 24924 (**C**). (**A**, **E)** adapted from Wick and Lehman (2013); (**B)** adapted from Gilmore (1919); (**C)** adapted from Wiman (1930); (**D)** adapted from Hatcher et al. (1907).

#### Figure S17

**"*Pentaceratops aquilonius*" (CMN 9813) shown to scale with c.f. *P. sternbergii* (AMNH 1625)**

Longrich (2014) proposed the new taxon *Pentaceratops aquilonius* (holotype CMN 9813, **A**) based in part on the suggestion that compared to c.f. *Pentaceratops sternbergii* (AMNH 1625, **B**), the parietal posterior bar is anteroposteriorly broad and only weakly embayed. This is based on inaccurate reconstruction of CMN 9813, mainly because of inappropriate scaling. This figure shows CMN 9813 is comparable to AMNH 1625 in the anteroposterior thickness of the posterior bar, and that CMN 9813 is too incomplete to infer size or shape of the median embayment. (**A**, **B**) Adapted from Longrich (2014). Scalebar equals 10 cm.

#### Figure S18

**Proposed reconfiguration of epiparietal numbering system in *Chasmosaurus* and related chasmosaurines**

The original epiparietal numbering systems of Holmes et al. (2001; **A**, **B**) and Sampson et al. (2010, **C**) are reconfigured (**G**-**I**) based on comparison to stratigraphically preceding chasmosaurines c.f. *Chasmosaurus russelli* (CMN 2280, **D**), and *C. belli* (ROM 843, **E**; YPM 2016, **F**). Most notably, locus ep1 develops from an spindle shaped epiparietal in c.f. *C. russelli* (**D**) to an elongate and anteriorly curving ridge in *C. belli* (**E**, **F**). In YPM 2016 the ep1 ridge bears 3-4 anteriorly projecting processes which are here interpreted to be homologous with the anteriorly projecting processes also in this position in the stratigraphically succeeding taxa *Vagaceratops* (*Chasmosaurus*) *irvinensis* (**G**, **H**) and *Kosmoceratops richardsoni* (**I**). Specimens not shown to scale.

#### Figure S19

**Angle of postorbital horn relative to maxillary tooth row in Chasmosaurinae sp. "*Agujaceratops mariscalensis*" TMM 43098-1 (A) and *Pentaceratops sternbergii* AMNH 6325 (B)**

In their rediagnosis of *Agujaceratops mariscalensis*, Forster et al. (1993) suggest "erect supraorbital horncores that attain an angle of 85° to the maxillary tooth row in adults" as a new autapomorphy, based on TMM 43098-1 (**A**). This figure shows that the postorbital horns of TMM 43098-1 are no more erect than those of *Pentaceratops sternbergii* holotype AMNH 6325 (**B**). The posterior curvature of the postorbital horns in TMM 43098-1 create the illusion of more erect horns. (**A**) adapted from Forster et al., (1993); (**B**) adapted from Osborn (1923).
