## Supplementary material for "Transitional evolutionary forms and stratigraphic trends in chasmosaurine ceratopsid dinosaurs : evidence from the Campanian of New Mexico": SUPP TXT2 - Phylogenetic anlaysis, characters

SUPPORTING INFORMATION 3

### PHYLOGENETIC ANALYSIS

Phylogenetic analysis was conducted using an adapted version of the character matrix from Mallon et al. (2014), which itself was adapted from Wick and Lehman (2013), Mallon et al. (2011), and originally, Sampson et al. (2010). The character matrix was edited in Mesquite 3.02 (Maddison and Maddison, 2015). Edits were made to 22 characters, including definitions, character states, and previous codings. Four new characters were added concerning the morphology of the parietal, making a total of 156 characters (see Supporting Information 2 for details of changes made and new characters). The taxon *Mojoceratops perifania* was recoded where necessary, but ultimately was removed from the matrix as it is a junior synonym of *Chasmosaurus russelli* (Maidment and Barrett, 2011). The two new taxa (Taxon A and Taxon B) were added to the matrix to make a total of 26 taxa. *Leptoceratops gracilis* was selected as the outgroup.

Analyses were rerun to investigate the effects of removing problematic fragmentary taxa. The first reanalysis removed *Bravoceratops polyphemus* and *Agujaceratops mariscalensis*, leaving 24 taxa. The second reanalysis further removed *Coahuilaceratops magnacuerna*, leaving 23 taxa.

The analyses were performed with parsimony using PAUP v. 4.0 b10 for Mac OSX (Swofford, 2002) within a heuristic search of 5,000 replicates using ACCTRAN optimization and tree bisection-reconnection swapping to produce a strict consensus tree, followed by a bootstrap analysis using a heuristic search of 5,000 replicates. Tree figures were prepared using FigTree 1.4.2 (Rambaut, 2014).

#### Phylogenetic character alterations

It has not been possible to review all character states for every taxon listed by Sampson et al. (2010) and Mallon et al. (2014). However, corrections were made in cases where we were familiar with the specimens (especially *Chasmosaurus*, *Pentaceratops*, and *Triceratops*). All corrections are noted below, and most concern parietal morphology.

#### Notes on specimens and data used for analysis

Additional specimens used to code character states are noted where appropriate.

##### *Bravoceratops*

The matrix has been amended to reflect the re-identification of the parietal median bar of *Bravoceratops polyphemus* (Wick and Lehman, 2013) as representing the anterior rather than posterior end (see main text). This results in characters 65, 68, 72, 76, 82-84, 95, 96, 98, 99, and 104-107 all being recoded as unknown (?).

##### *Vagaceratops* and *Kosmoceratops*

The character matrix for *Vagaceratops* (*Chasmosaurus*) *irvinensis* and *Kosmoceratops* *richardsoni* (Holmes et al., 2001; Sampson et al., 2010; Loewen et al., 2013) has been altered to reflect reinterpretation of epiparietals and episquamosals. This is discussed in the main text, and results in changes for character states 88-90, 92, 96, 97, 102, and 103.

#### Notes & alterations to existing characters

##### Character 7

"Premaxilla, narial strut orientation (Dodson et al., 2004, character 6)"

0: 'rostrally inclined'

1: 'caudally inclined'

*Coahuilaceratops* recoded from (1) to (?) as the premaxillary strut is heavily damaged and might not be caudally inclined. *Chasmosaurus belli* recoded from (0) to (0&1) as YPM 2016 has a caudally inclined narial strut.

##### Character 60

"Squamosal, length relative to parietal (Holmes et al., 2001, character 19, modified)"

0: 'squamosal much shorter than parietal, large portion of posterolateral frill made up of parietal'

1: 'squamosal slightly shorter than parietal, pmn;y posterolaterl-most [sic] margin of frill formed by the parietal'

2: 'squamosal and parietal equal in length'

*Pentaceratops sternbergii* is recoded from (1) to (1&2). *P. sternbergii* specimens AMNH 1624 and 1625 exhibit a squamosal that is the same length as the parietal, although in other specimens (UKVP 16100, MNA Pl.1747) the squamosal is slightly shorter (state 1). *Anchiceratops* is recoded from (2) to (1&2), as some specimens have squamosals that are equal in length to the parietal (e.g. ROM 802), whereas others the squamosal is shorter (e.g. AMNH 5251). *Ojoceratops* is originally coded as (2), however no squamosal has been collected with an associated parietal so that this cannot be asserted (Sullivan and Lucas, 2010). *Ojoceratops* is therefore recoded as (?).

##### Character 61

Squamosal forms part of posterior margin of frill (new)

0: 'present '

1: 'absent '

Ojoceratops changed from (1) to (?) for the same reason as the above character 60.

##### Character 66

"Parietal, concave median embayment on caudal margin (new)"

0: 'absent '

1: 'present '

*Torosaurus latus* and *Triceratops horridus* both changed from state (0) to (0&1). YPM 1831 (holotype of *Torosaurus* "*gladius*"=*latus*) has a clear embayment on the midline of the posterior parietal, as does *Triceratops horridus* specimen AMNH 5116. However, other specimens of *Torosaurus latus* (e.g. MOR 1122) and *Triceratops horridus* (e.g. MOR 1120) lack a medial embayment (state 0). A possible very weak embayment is present in isolated parietals MOR 2946 and MOR 6647, which probably both pertain to *Triceratops prorsus*; MOR 2946 exhibits a stepped margin to the incipient fenestra on the ventral side (only seen in *T. prorsus*; Scannella and Horner, 2010); and MOR 6647 is from the upper third of the Hell Creek Formation (pers. obs.), from which only *T.* *prorsus* remains are known (Scannella et al., 2014). However, given that the embayment in these specimens is not very pronounced, I am going to leave *Triceratops prorsus* as state (0).

*Chasmosaurus belli* is recoded from (0) to (1). It is noted that in the original coding, *C. belli* is coded as lacking an embayment in character 66 (state 0), but in the following character (67) is coded as character state (1): 'shallow, entire transverse bar is a V-shaped embayment' (ie. that it possesses an embayment). The lateral rami of the parietal posterior bar are inclined in all specimens attributed to *C. belli*, meeting at the midline as a shallowly concave embayment (ie. character 66, state 1). It is noted here that the angle at which the lateral rami meet is almost flat in YPM 2016, being intermediate between the condition seen in other *C. belli*, and that of *Vagaceratops* (*Chasmosaurus*) *irvinensis*.

Although *Anchiceratops* is coded as lacking a medial embayment, if Lehman (1993, 1998) is correct then this is because the embayment has closed, rather than it never having existed in the ancestor. Furthermore, in all *Anchiceratops* specimens, the midline of the parietal posterior bar (point 2 in my morphometric analysis) is notably more anterior than the posteriormost part of the parietal (excluding epiparietals); i.e. it is embayed. This narrow notch-like embayment occurs between the epiparietals. Why this is not therefore coded as possession of an embayment is not clear. *Pachyrhinosaurus lakustai* and *Centrosaurus apertus* are both coded for the possession of a median embayment, yet it is very shallow, and in at least two specimens of *P. lakustai* (TMP 1987.55.141 and 1988.55.46; Currie et al., 2008) it is extremely shallow to absent.

##### Character 67

"Parietal, shape of concave median embayment (Forster, 1990, character 83, modified) PROBLEM WITH NUMBERS ON SHEET"

0: 'shallow, restricted to center of margin '

1: 'shallow, entire transverse bar is a V-shaped embayment '

2: 'notch-like, restricted to center of margin'

As with character 66, *Torosaurus latus* (YPM 1831) and *Triceratops horridus* (AMNH 5116) both exhibit a shallow median embayment and hence these taxa are recoded from(?) to (0). Note that the "PROBLEM.." part of the description was in the original nexus file from Mallon et al. (2014).

##### Character 68

"Parietal, location of caudalmost point of caudal ramus (Holmes et al., 2001, character 18, modified)"

0: 'on midline '

1: 'between midline and lateral-most corner '

2: 'at lateral-most corner adjacent to squamosal'

*Pentaceratops* has been recoded from state (2) to (1&2). AMNH 1625 is like *Utahceratops* (coded as 1) in that the caudalmost part of the parietal posterior bar is not adjacent to the squamosal. It could be argued that other *Pentaceratops* specimens coded as state 2 (MNA Pl.1747 and UKVP 16100) are in fact intermediate between states 1 and 2, and unlike the condition in *Chasmosaurus belli*, which is more definitively coded as state 2. *Triceratops horridus* has also been recoded from (0) to (0&1) as the mounted exhibit specimen AMNH 5116 clearly exhibits a central embayment in the parietal (hence character state 1; the midline is embayed so cannot be state 0).

##### Character 74

"Parietal, rim on medial margin of dorsotemporal fenestra (Forster, 1990, character 86). CORRECTED."

0: 'absent '

1: 'present, well-defined, laterally projecting rim defines medial margin of fenestra'

*Mojoceratops*, *Chasmosaurus* *russelli* and *C. belli* are recoded from (0) to (1). All specimens attributed to these taxa have a rim on the medial margin of the dorsotemporal fenestra.

Both species of *Torosaurus* and of *Triceratops* were originally coded as (0). This is partly understandable as the dorsotemporal fenestrae are not as prominent in *Torosaurus* and *Triceratops*, and the ridge is weak in large (probably mature) specimens. Nevertheless, a rim on the dorsotemporal fenestra is visible in *Torosaurus latus* specimen MOR 1122; in *Triceratops* *horridus* specimens USNM 2100 and AMNH 5116; in immature specimens of *Triceratops prorsus* held at MOR, and in the holotype (YPM 1822; see Hatcher et al., 1907; pl. 35). This region of the parietal is fragmentary in *Torosaurus* *utahensis*, but appears to preserve a rim (Scannella, pers. comm.). Hence *Torosaurus* and *Triceratops* are recoded from (0) to (1).

##### Character 76

"Parietal, rostrocaudal thickness of transverse bar at narrowest point (Holmes et al., 2001, character 22)"

0: 'narrow and straplike, less than 10% total parietal length'.

1: 'broad, 20% or more of total parietal length.'

Despite being incomplete, Taxon A and Taxon B are coded for this character based on reconstruction of the parietal (see figures in main text), and the assertion that the rostrocaudal thickness of the narrowest point of the lateral ramus of Taxon B cannot be 10% or less of total parietal length without the parietal being unusually elongate.

##### Character 77

"Parietal, relative rostrocaudal depth of broad transverse bar (new)"

0: 'subequal medial to lateral'.

1: 'tapering so that the narrowest point occurs medially'.

This was recoded as (0&1) for *Pentaceratops sternbergii* (previously 1) because AMNH 1625 records state 0 as it does not notably taper mediolaterally. Tapering is noted, however, in UKVP 10161, and MNA Pl.1747 (both state 1). This has also been recoded for *Utahceratops*, which was coded as (1) and is recoded as (0). Despite being incomplete, the parietal of *Utahceratops* does not narrow notably medially. Indeed, in the reconstruction offered by Sampson et al. (2010), if anything the parietal is rostrocaudally deeper medially rather than laterally (although only very slightly).

##### Character 78

"Parietal, cross-sectional shape of median bar (Holmes et al., 2001, character 24, modified)"

0: 'diamond-shaped, tapers laterally'

1: 'rectangular or subrectangular, margin facing parietal fenestrae thick and oriented sub-perpendicular to parietal surface '

2: 'round to lenticular'

3: 'v-shaped, opening ventrally'

*Pentaceratops* altered to state 0 and 2 (previously only 2) as the MNA Pl.1747 specimen of *Pentaceratops* possesses a lenticular cross section with slight lateral flanges, similarly seen in Taxon A and Taxon B.

This character state is poorly defined. None of the defined character states adequately describe the condition seen in Taxon A, Taxon B*,* or indeed Taxon C; furthermore, the cross section of the median bar changes along its length and the character as defined makes no claim as to where it should be sampled. Taxon A and Taxon B are therefore each coded as (0&2).

##### Character 86

"Episquamosal, location of largest/longest example (new)"

0: 'episquamosals subequal in size '

1: 'rostralmost episquamosal by far the largest '

2: 'caudalmost episquamosal by far the largest'

There is a serious issue with this character in that the size of the anteriormost (rostralmost) episquamosal (aes) can often appear large because it is mounted on an anteroventral extension of the squamosal, and because (in *Triceratops* at least; Forster, 1996) this es is one of the first to fuse to the squamosal, and can therefore be difficult to delineate its actual size. The aes is often more pointed than other episquamosals (es), but is not obviously by far the largest in size (it is not clear if this means basal width, total area, etc).

A review of *Chasmosaurus belli* and *C. russelli* specimens does not reveal any specimens which unequivocally bear an aes in this position that is "by far the largest". Illustrations of *C. russelli* specimen NMC 2280 (Godfrey and Holmes, 1995) show the aes is roughly the same size as many other es (especially on the left squamosal). *C. russelli* specimen TMP 81.19.175 (Godfrey and Holmes, 1995) bears an aes that is slightly larger than the next most anterior es, but lacks the posteriormost es for comparison and so cannot be coded. *C. belli* specimen ROM 843 bears an aes that is larger than the next most anterior es, but comparable to some of the more posterior es. CMN 2245 bears an episquamosal that is well fused to the anteroventral projection such that it is not possible to tell how big it is based on photographs; regardless, the posterior es are quite large and it is not likely that the aes was by far the largest in this specimen. AMNH 5402 does not bear an aes that is any larger than other es positions. Based on these observations, *C. belli* and *C. russelli* are coded as (0).

The holotype of "*Mojoceratops*" *perfania* (TMP 1983.25.1) bears no obviously fused episquamosals and so should be coded as (?). Referred specimen AMNH 5656 has episquamosals that are of similar size along the squamosal, and so would be coded as (0) (although it is noted that this specimen is very small, and thus likely represents a juvenile, which may make its morphology unreliable).

*Bravoceratops* is recoded from (1) to (?). In their figure 8, Wick and Lehman (2013) illustrate the anterior end of the squamosal, which shows that the aes is actually small relative to other es. It appears large as it is mounted on the anteroventral extension of the squamosal. Although fragmentary, the squamosal of *Bravoceratops* bears six observable es, all of which are similar in size (ie. state 0). However, the posterior end of the squamosal is missing, such that it is not possible to exclude character state (2). As such, *Bravoceratops* must be coded as (?).

In contrast, the posteriormost episquamosal is often notably larger than other episquamosals, as recorded in the original dataset.

##### Character 88

Episquamosal locus S1 shape (new)

0: 'small and crescentic'

1: 'low raised D-shaped process'

2: 'well developed larger triangular process'

3: 'elongate hook'

Reconfiguration of the frill epiossifications of *Kosmoceratops* changes its state from (3) to (2).

##### Character 89

"Episquamosal locus S2 shape (new)"

0: 'small and crescentic'

1: 'low raised D-shaped process'

2: 'well developed larger triangular process'

*Vagaceratops* (*Chasmosaurus*) *irvinensis* is coded as (2), but this is inaccurate. The holotype (NMC 41357; Holmes et al., 2001) bears no episquamosals that can be considered as "a well developed larger triangular process" and should therefore be coded as (1). Referred specimen TMP 87.45.1 has some more triangular projections of the squamosal margin visible in photographs (Holmes et al., 2001), but these are in the middle of the squamosal, not in locus s2. Finally, *Vagaceratops* is originally coded as (0) for character 86 (Episquamosal, location of largest/longest example), suggesting that its episquamosals are all of similar size, which would contradict a coding of (2). *Vagaceratops* is here recoded from (2) to (1).

Reconfiguration of the frill epiossifications of *Kosmoceratops* changes its state from (2) to (1). Note that in the holotype of *Kosmoceratops* UMNH VP 17000, the reconfigured es2 (es3 in the original configuration) is a D-shape, whereas it is more pointed in referred specimen UMNH VP 16878 (Loewen et al., 2013).

##### Character 90

"Episquamosal locus S2 size relative to other episquamosals (new)"

0: 'subequal'

1: 'second only to S1 in size, larger than S3'

Reconfiguration of the frill epiossifications (es3 is renumbered here as es2) of *Kosmoceratops* changes its state from (1) to (0).

##### Character 91

"Marginal ossification crossing squamosal-parietal contact (Dodson et al., 2004, character 43)"

0: 'absent'

1: 'present'

This character refers to the presence/absence of a frill epiossification that straddles the squamosal-parietal contact (referred to hereafter as an eps).

The character is of dubious utility for two reasons. Firstly, in many specimens epiossifications are not fused on to the frill border such that lack of a visible eps does not mean that one was absent in life. The only way that state (0) can be unequivocally asserted is if a specimen has epiossifications fused to either the squamosal or parietal such that there is no available space for an eps. Secondly, even if it is possible to assess the absence of an eps in one specimen, possession of an eps is observed to be very variable individually in some taxa. For example, the most well-known chasmosaurine, *Triceratops* typically exhibits 5-7 epiparietals (Scannella et al., 2014). In specimens with 5 epiparietals, an eps (or space for one) is typically present. In specimens with 7 epiparietals, an eps is typically absent (i.e. in some specimens the eps might "jump" on to the parietal).

In light of these issues, this character is miscoded for many taxa, so I have made a number of revisions. *Pentaceratops* *sternbergii* was previously coded as (0) but is here recoded as (0&1). Only *P. sternbergii* specimen AMNH 1624 can be reliably coded as (0) as it preserves fused epiossifications which do not straddle the parietal-squamosal contact, nor is there a space for an eps. However, *P.* *sternbergii* frill AMNH 1625 (not included in the observed specimen list of Sampson et al., 2010) clearly possesses a triangular epiossification that straddles the parietal-squamosal contact. *Nedoceratops* is recoded from (1) to (?) as the parietal-squamosal contact is only preserved on the right side of the skull, and even here it is incomplete and may or may not preserve an eps (Farke, 2011).

*Eotriceratops* and *Torosaurus* *latus* are treated inconsistently. *Eotriceratops* was originally coded as (1), which is probably true as there seems to be a space at the distal end of the squamosal where an eps might have occurred (and is inferred to have been by Wu et al., 2007). The parietal of *Eotriceratops* is unknown, but presumably on the basis of this space, and the logic outlined in the above paragraph, Sampson et al. (2010) code an eps as present. However, *Torosaurus latus* is coded as (0), despite possessing a notable gap at the parietal-squamosal contact in MOR 1122 (which was studied by Sampson et al., 2010). Furthermore, *Torosaurus latus* specimen ANSP 15192 exhibits an eps (Scannella and Horner, 2010), but this is not noted by Sampson et al. (2010; who also studied this specimen). Therefore I have recoded *Torosaurus latus* as (0&1). As mentioned above, specimens of *Triceratops* exhibit both conditions and should therefore be recoded from (1) to (0&1).

Reconfiguration of the frill epiossifications of *Kosmoceratops* and *Vagaceratops* does not change the character state, but what was previously considered as es1 is here coded as an ep3 that is possibly partly borne on the squamosal as an eps.

##### Character 92

"Shape of marginal ossification crossing squamosal-parietal contact (new)"

0: 'small and crescentic'

1: 'present strongly recurved process or gnarled process'

2: 'well developed triangular process sometimes with a small peak'

Note that no taxa are coded as state (0), hence this character state is redundant. More importantly, the description of the morphology of eps is inconsistent with morphological description of other epiossifications. For example:

Character 102. "Epiparietal locus P3 shape (new)"

0: 'low D-shaped process '

1: 'elongate flattened process or spike '

2: 'strongly recurved triangular or recurved low gnarled triangular process'

3: 'well developed triangular process'

4: 'elongate low process sometimes with a small peak'

This is significant as the current definition of character 92 conflates "well-developed triangular process" with the "elongate low process" as a single character. It is suggested here that if the eps were coded using the morphological definitions of character 102, some taxa would not be lumped together as state (2); for example, the eps of *Anchiceratops* and *Pentaceratops* (see below) is triangular in shape and unlike the spindle-shaped eps seen in *Triceratops* and *Arrhinoceratops* (which is basically the same morphology as most other epiossifications of the frill). Therefore we have revised the character state definitions of character 92 as follows:

##### Character 92 (revised)

"Shape of marginal ossification (eps) crossing squamosal-parietal contact (new)"

0: 'small and crescentic'

1: 'present strongly recurved process or gnarled process'

2: 'well developed triangular process'

3: 'elongate base, spindle-shaped low process sometimes with a small peak'

4: 'elongate hook'

State (0) is conserved as presumably this permits comparison with other datasets.

*Albertaceratops* and *Centrosaurus* were both coded as (0) for character state 91 (i.e. they do not possess an eps), but were coded (0) for character 92. This is in error. I have recoded *Albertaceratops* and *Centrosaurus* as (?) for character 92. *Pentaceratops* *sternbergii* was previously coded as (?) but is here recoded as (2). *P.* *sternbergii* frill AMNH 1625 (not included in the observed specimen list of Sampson et al., 2010) clearly possesses a triangular epiossification that straddles the parietal-squamosal contact. *Anchiceratops* remains as (2). *Eotriceratops* was previously coded as (2), i.e. the same as other triceratopsins. However, although the presence of an eps is inferred (see Wu et al., 2007), the actual eps is not present, hence its morphology cannot be known. *Eotriceratops* is therefore recoded as (?). *Arrhinoceratops* and all species of *Triceratops* and *Torosaurus* are recoded from (2) to (3).

A further problem exists in that the identity of the epiossification at position eps might vary, and in at least some cases, character 92 might be coding for the morphology of ep3, which is already covered by character 103. This is probably the case with *P. sternbergii* specimen AMNH 1625 as it possesses an ep1 within the parietal embayment, an ep2 approximately halfway along the lateral rami, and an epiossification positioned as eps, which should probably be considered as ep3. Unfortunately no other specimens of *P. sternbergii* exist which preserve three epiossifications contacting the parietal such that comparisons can be made. Further discussion of the identity of ep3/eps can be found in the discussion in main text. We have chosen to leave character 92 in the analysis (in its revised form), but acknowledge a case can be made for its removal.

Reconfiguration of the frill epiossifications of *Kosmoceratops* changes the character state from (1) to the new state (4). Note that the laterally curving hook-shaped epiossification previously considered as es1 is here coded as an ep3 that is possibly partly borne on the squamosal as an eps.

Reconfiguration of the frill epiossifications of *Vagaceratops* changes the character state from (1) to state (2).

##### Character 96

"Epiparietal locus DPP1 (Sampson, 1995, character 14, modified)"

0: 'absent '

1: 'present '

This character refers to the possession of epiparietal locus "DPP1" but is not referred to as this in Sampson (1995) from which it is supposedly modified (a paper concerned with the description of new centrosaurine taxa). Character 14 of Sampson (1995, p. 760) is defined as "Posterior parietal hooks (Process 1): (0) absent; (1) present, abbreviated; (2) present, well-developed". Subsequent characters follow the same format for epiparietals 2 to 6; for example character 15 is defined as "Process 2 on parietal margin: (0) absent; (1) small; (2) large, medially-directed horn. Therefore it is established here that character 14 of Sampson (1995) refers to the possession and (if present) morphology of an epiparietal (parietal process) at position 1 (all in one character).

The current character 96 only has two character states (0) absent, and (1) present, hence it is not fully comparable to character 14 of Sampson (1995). If character 96 is simply supposed to record the presence / absence of epiparietal 1 (ep1), then this is understandable, however, epiparietals 2 and 3 are not afforded their own presence-absence character (although there is less variability in ep2 and ep3). Curiously, *Utahceratops* and *Pentaceratops* are the only chasmosaurines coded as state (1) for character 96, which might therefore suggest that the ep1 locus is missing from other chasmosaurine taxa. This might be understandable, as *Chasmosaurus belli* exhibits an elongate ridge in the position of ep1, so it could be arguable that the ep1 locus is therefore absent in this taxon (I would disagree; I think the ridge is an laterally expanded ep1). However *C. belli*, and indeed all chasmosaurines (for which the posterior bar of the parietal is preserved) are coded for morphology of ep1 in character 97, which therefore shows that according to the original coding, they indeed possess the ep1 locus (we agree). This confusing issue would appear to be solved by either coding all chasmosaurines as state (1) for character 96, or deleting / altering character 96.

Note that *Pachyrhinosaurus* *lakustai* (Currie et al., 2008) is coded by Sampson et al. 2010 and Mallon (2014) as state (1), ie. that it possesses an ep1. However, in the original description, the parietal of *P.* *lakustai* is shown as lacking ep1. Therefore we have recoded *P. lakustai* as state (0).

##### Character 96 (revised)

"Epiparietal locus ep1, position (New configuration)"

0: absent

1: present; locus ep1 is positioned at the lateral edge of the parietal embayment or the space usually occupied by an ep0, but not within this space; this is usually where there is a change of angle in the posterior bar.

2: present, locus ep1 is elongate, and extends laterally from the midline (or near the midline) over 50% or more of each lateral ramus of the parietal posterior bar.

3: present, locus ep1 is limited to the central third of the posterior bar, such that the medialmost edge of locus ep1 immediately abuts the midline (or nearly so).

*Leptoceratops*, *Protoceratops*, *Albertaceratops* and *Pachyrhinosaurus* are coded as (0). *Centrosaurus* is coded as (3).

*Chasmosaurus russelli*, *Mojoceratops perfania*, and *Agujaceratops mariscalensis* all exhibit character state (1).

*Chasmosaurus belli*, *Vagaceratops* (*Chasmosaurus*) *irvinensis* and *Kosmoceratops* *richardsoni* exhibit character state (2). This is part of the reconfiguration of epiparietal classification within this study. Most specimens of *C. belli* exhibit a simple anteriorly recurved elongate ridge in the ep1 locus position, however one specimen (YPM 2016) has what appears to be an array of protuberances developed along this ridge, but is otherwise like other specimens of *C. belli*. These protuberances are here suggested to develop into the prominent processes seen on the parietal of *Vagaceratops* (*Chasmosaurus*) *irvinensis*, and further developed into more elongate spikes in *Kosmoceratops*.

*Utahceratops*, *Pentaceratops* *sternbergii*, *Anchiceratops*, Taxon A, and Taxon B all exhibit character state (3).

*Arrhinoceratops*, *Ojoceratops* (based on the parietal described by Farke and Williamson, 2006; also used by Sampson et al. 2010), *Torosaurus* *latus*, and both species of *Triceratops* are coded as (1).

##### Character 97

"Epiparietal, locus P1 shape (Sampson, 1995, character 15, modified)"

0: 'low D-shaped process'

1: 'elongate flattened process or spike'

2: 'strongly recurved triangular or recurved low gnarled triangular process'

3: 'well developed triangular process'

4: 'elongate low process sometimes with a small peak'

*Albertaceratops* (Ryan, 2007) is coded by Sampson et al. 2010 and Mallon (2014) as state (1), ie. that it possesses an ep1. However, in the original description, the parietal of *Albertaceratops* is shown as lacking ep1, and is coded as such in the data matrix (Ryan, 2007). *Albertaeratops* is also coded as lacking an ep1 in character 96. Here we have therefore coded *Albertaceratops* as (?) for character 96 (it should really be "-", but in this analysis absent characters are coded as ?; see note). Similarly, epiparietal 1 is not present in *Pachyrhinosaurus* *lakustai* (Currie et al., 2008), hence I have recoded this from (1) to (?). Note that no taxa in the current matrix possess character state (0); presumably this is a result of culling other taxa from the matrix in a dataset prior to Sampson et al. (2010).

Character 97 has problems as most chasmosaurine frill epiossifications broaden and become less pointed through ontogeny; this has been demonstrated unequivocally in *Triceratops* (Horner and Goodwin, 2006; 2008) and is observable in other chasmosaurines (e.g. Lehman, 1990; Godfrey and Holmes, 1995; Mallon et al., 2014). However, in *Anchiceratops* and possibly *Pentaceratops*, parietal epiossifications remain triangular and pointed even in the largest and presumably oldest individuals (Mallon et al., 2011; see specimen descriptions in supporting information 1). Moreover, it is not very clear what the difference is between character state 1 and 4 (for example). By reference to the codings (*Centrosaurus* is the only taxon coded as state 1) it can be seen that character state (1) refers to anteroposterior elongation, whereas state (4) refers to the broadening of the base of the epiossifications (in the case of the parietal this would usually be lateral broadening, but on the squamosal, usually anteroposterior).

These issues are reflected in the revised character 97:

##### Character 97 (revised)

"Epiparietal, locus P1 morphology (revised)"

0: 'low D-shaped process '

1: 'anteroposteriorly elongate flattened process or spike '

2: 'strongly recurved triangular or recurved low gnarled triangular process'

3: 'well developed triangular process'

4: 'elongate base, spindle-shaped low process sometimes with a small peak'

5: 'very elongate and differentiated into an array of (typically 3) protuberances'

Note the addition of state (5): *Vagaceratops* *irvinensis* and *Kosmoceratops* *richardsoni* both exhibit an array of anteriorly projecting protuberances along the posterior bar of the parietal; this is coded as state (5). One specimen of *Chasmosaurus belli* (YPM 2016) has what appears to be an incipient development of this character (the protuberances are less pronounced, but clearly present). Other specimens of *C. belli* (e.g. CMN 491, ROM 843, BMNH 4948) exhibit an elongated ridge in the position of ep1, but it is an undifferentiated single ridge. Therefore *C. belli* is coded as (2&5). *Mojoceratops* is recoded as (2).

##### Character 98

"Epiparietal, locus P1 orientation (Sampson, 1995, character 15, modified)"

0: 'caudally, epiparietal oriented in the plane of the frill '

1: 'directed rostrodorsally '

2: 'P1 occurs on dorsal surface of parietal'

Again, both *Albertaceratops* and *Pachyrhinosaurus* lack an epiparietal 1, so should be coded as (?) instead of the original coding of (0). Also, since the parietal of *Bravoceratops* was misidentified (see main text), then it is now coded as (?), leaving only one taxon (*Anchiceratops*) that would exhibit character state (2) (which was added by Wick and Lehman, 2013, and maintained by Mallon et al., 2014). This makes character state (2) redundant. I am therefore reverting the character back to as it was in Sampson et al. (2010).

##### Character 98 (revised)

"Epiparietal, locus P1 orientation (Sampson, 1995, character 15, modified)"

0: 'caudally, epiparietal oriented in the plane of the frill '

1: 'directed rostrodorsally '

Reconfiguration of the frill epiossifications of *Kosmoceratops* and *Vagaceratops* does not result in a change in character state, although what was previously considered as ep1, ep2, and ep3 by (Sampson et al., 2010; Loewen et al., 2013) are here considered as an array of processes hosted on locus ep1.

##### Character 99

" Epiparietal, locus P1 curvature (Sampson, 1995, character 15, modified)"

0: 'straight '

1: 'laterally curved '

2: 'medially curved'

3: 'dorsally curved'

Reconfiguration of the frill epiossifications of *Kosmoceratops* and *Vagaceratops* does not result in a change in character state, although what was previously considered as ep1, ep2, and ep3 by (Sampson et al., 2010; Loewen et al., 2013) is here considered as an array of processes hosted on ep1.

##### Character 100

" Epiparietal, locus P2 shape (Sampson, 1995, character 16, modified)"

0: 'low raised D-shaped process'

1: 'elongate spike'

2: 'strongly recurved triangular or recurved low gnarled triangular process'

3: 'well developed triangular process'

4: 'elongate low process sometimes with a small peak'

revised for clarity:

##### Character 100 (revised)

" Epiparietal, locus P2 shape (Sampson, 1995, character 16, modified)"

0: 'low D-shaped process'

1: 'anteroposteriorly elongate flattened process or spike'

2: 'strongly recurved triangular or recurved low gnarled triangular process'

3: 'well developed triangular process'

4: 'elongate base, spindle-shaped low process sometimes with a small peak'

Reconfiguration of the frill epiossifications of *Kosmoceratops* and *Vagaceratops* does not result in a change in character state, although what was previously considered as eps by (Sampson et al., 2010; Loewen et al., 2013) is here considered as an ep2.

##### Character 101

" Epiparietal, locus P2 curvature (Sampson, 1995, character 16, modified)"

0: 'straight '

1: 'medially or laterally curved in the plane of the frill'

2: 'recurved onto dorsal surface of frill'

Reconfiguration of the frill epiossifications of *Kosmoceratops* and *Vagaceratops* does not result in a change in character state, although what was previously considered as eps by (Sampson et al., 2010; Loewen et al., 2013) is here considered as an ep2.

##### Character 102

"Epiparietal locus P3 shape (new)"

0: 'low raised D-shaped process'

1: 'elongate spike'

2: 'strongly recurved triangular or recurved low gnarled triangular process'

3: 'well developed triangular process'

4: 'elongate low process sometimes with a small peak'

revised for clarity and addition of a new state:

##### Character 102 (revised)

"Epiparietal locus P3 shape (new)"

0: 'low D-shaped process'

1: 'anteroposteriorly elongate spike'

2: 'strongly recurved triangular or recurved low gnarled triangular process'

3: 'well developed triangular process'

4: 'elongate base, spindle-shaped low process sometimes with a small peak'

5: 'elongate hook'

Reconfiguration of the frill epiossifications of *Kosmoceratops* changes the character state from (1) to the new state (5). Note that the laterally curving hook-shaped epiossification previously considered as es1 is here coded as an ep3 that is possibly partly borne on the squamosal as an eps (see characters 91 and 92) .

Reconfiguration of the frill epiossifications of *Vagaceratops* changes the character state from (2) to state (3).

##### Character 103

"Epiparietal, locus P3 orientation (new)"

0: 'caudally, epiparietal oriented in the plane of the frill'

1: 'directed rostrodorsally'

revised to:

##### Character 103 (revised)

"Epiparietal, locus P3 orientation (new)"

0: 'caudally, epiparietal oriented in the plane of the frill'

1: 'directed rostrodorsally'

2: 'directed laterally'

Reconfiguration of the frill epiossifications of *Kosmoceratops* changes the character state from (1) to the new state (2). Note that the laterally curving hook-shaped epiossification previously considered as es1 is here coded as an ep3 that is possibly partly borne on the squamosal as an eps (see characters 91 and 92) .

Reconfiguration of the frill epiossifications of *Vagaceratops* changes the character state from (1) to state (0).

*Mojoceratops* was originally coded as (0), but in the two specimens that preserve ep3 (holotype TMP 1983.25.1, AMNH 5656) it is clearly oriented laterally to caudolaterally, and is hence recoded as (2). Although fragmentary, part of the parietal of *Agujaceratops* is preserved which probably represents ep3 (UTEP P.37.7.065; Lehman, 1989), and may indicate that it is directed laterally. However, due to the fragmentary nature of this specimen, here we have chosen to leave *Agujaceratops* coded as (?).

*Chasmosaurus russelli* is recoded from (0) to (0&2). Although the drawing in Godfrey and Holmes (1995) shows the ep3 as caudolaterally directed on both sides, in the fossil the right ep3 is laterally directed. This does not seem to be a result of significant distortion. *C. belli* is recoded from (0) to (2). Ep3 is laterally or caudolaterally positioned in YPM 2016, ROM 843, and CMN 2245.

#### New characters

The posterior border of the parietal is the single most diagnostic element in chasmosaurines. These new characters help define differences in parietal morphology that are not covered by the character matrix of Sampson et al. (2010) or Mallon et al. (2014).

##### Character 153 (new)

"Parietal, position of the maximum constriction of the median bar"

0: 'within the anterior 2/3 of the parietal fenestra'

1: 'within the posteriormost 1/3 of the parietal fenestra'

The median bar of the parietal narrows (constricts) to give a "waisted" shape. Specimens vary conspicuously as to the anteroposterior position of the maximum point of constriction. Centrosaurines, *Kosmoceratops*, *Vagaceratops*, and species of *Chasmosaurus* have a maximum constriction point in the posteriormost third of the median bar, often very close to where the median bar meets the posterior bar. By contrast, *Pentaceratops*, *Anchiceratops*, *Arrhinoceratops* and *Torosaurus* exhibit a point of maximum constriction that is roughly at the midpoint of the median bar (as defined by the borders of the parietal fenestrae).

##### Character 154 (new)

"Epiparietal, locus P1 orientation, angle of the long axis of ep1 relative to the long axis of the parietal (ORDERED)"

0: 0°-30°

1: 30°-60°

2: 60°-90°

3: 90°+

A major difference between specimens is the angle of the long axis of ep1. The character is measured by drawing a line through the lateral and medial edges of the epiparietal and measuring the angle it makes with the long axis of the parietal (measured along the median bar). This character closely matches the angle formed by the medial meeting of the lateral rami of the posterior bar, but is more easily measurable.

##### Character 155 (new)

"Parietal, thickness of transverse/posterior bar at thickest point (ORDERED)"

0: narrow and strap-like (1-10 % of maximum parietal width in posterior half)

1: medium thickness (11-15 % of maximum parietal width in posterior half)

2: thick (15-20 % of maximum parietal width in posterior half)

3: very thick (>20% of maximum parietal width in posterior half)

"Maximum parietal width in posterior half " is used as a proxy for size and is chosen over parietal length (as in character 76) as parietal length is very difficult to measure in most specimens as the anterior end is very fragile. This character helps differentiate between taxa that have a uniformly anteroposteriorly thin posterior bar, and those that have a thick posterior bar. Specimens where only one half of the posterior bar is fully preserved have the preserved side measured, then doubled. *Anchiceratops* specimen CMN 8535 was not measured for this character as the lateral width of the parietal is not possible to measure as the parietal-squamosal contact is difficult to discern. *Utahceratops* was coded from the reconstruction in Sampson et al. (2010), which was considered as accurate enough for this measurement.

##### Character 156 (new)

"Parietal fenestrae: shape (ORDERED character)"

0: angular

1: subangular

2: subrounded

3: rounded

Shape definition based on the same visual criteria as sediment grain roundedness.
