## Supplementary figures and images for "Transitional evolutionary forms and stratigraphic trends in chasmosaurine ceratopsid dinosaurs : evidence from the Campanian of New Mexico"

### Figure S1

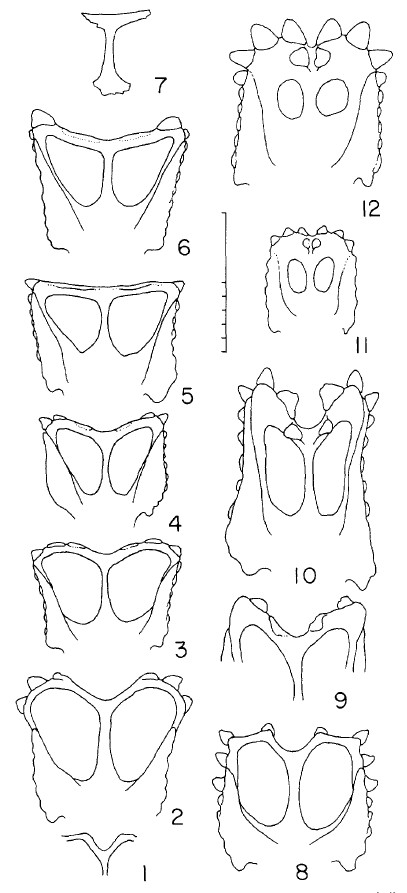

### Figure S2

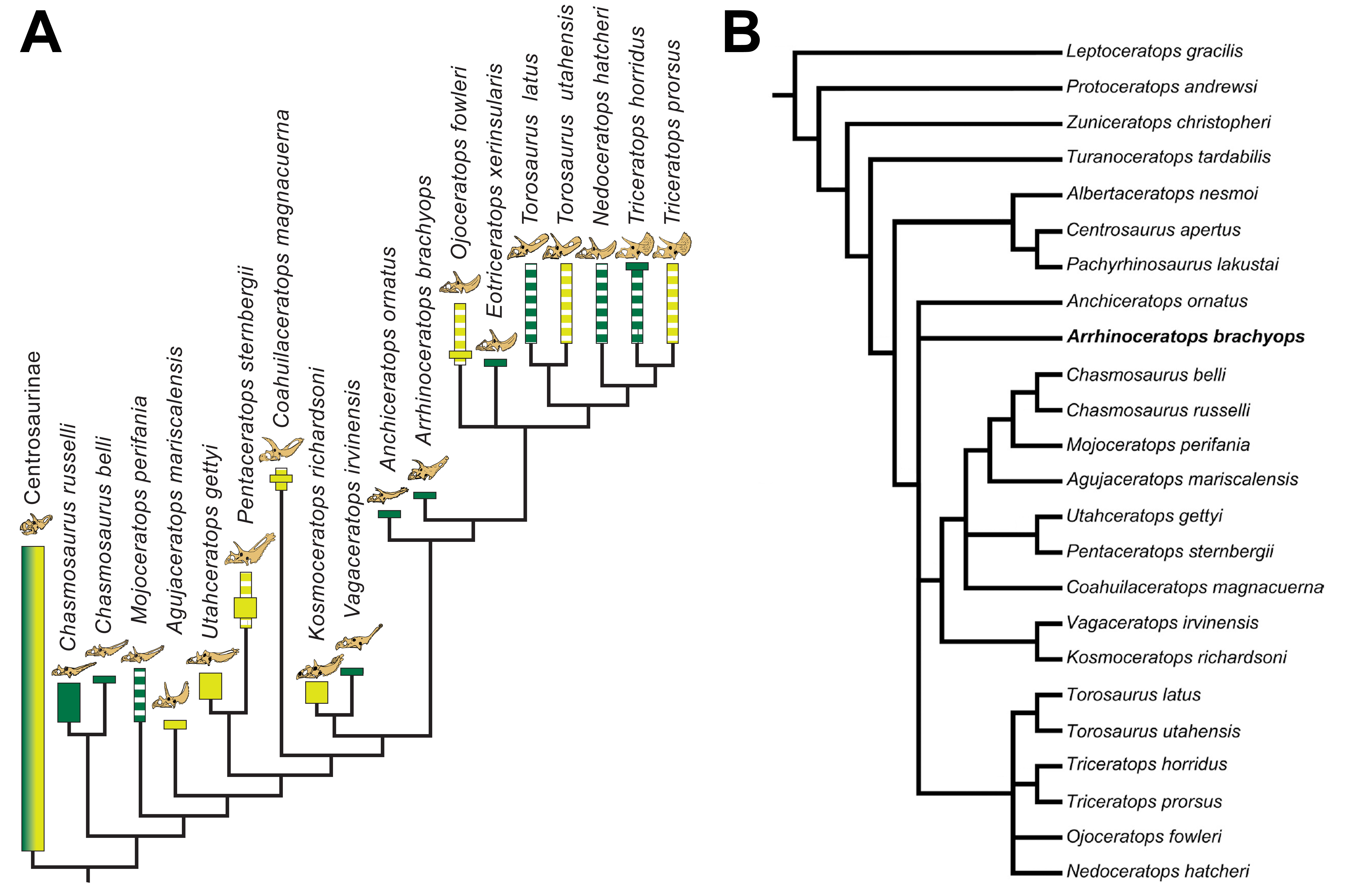

### Figure S3

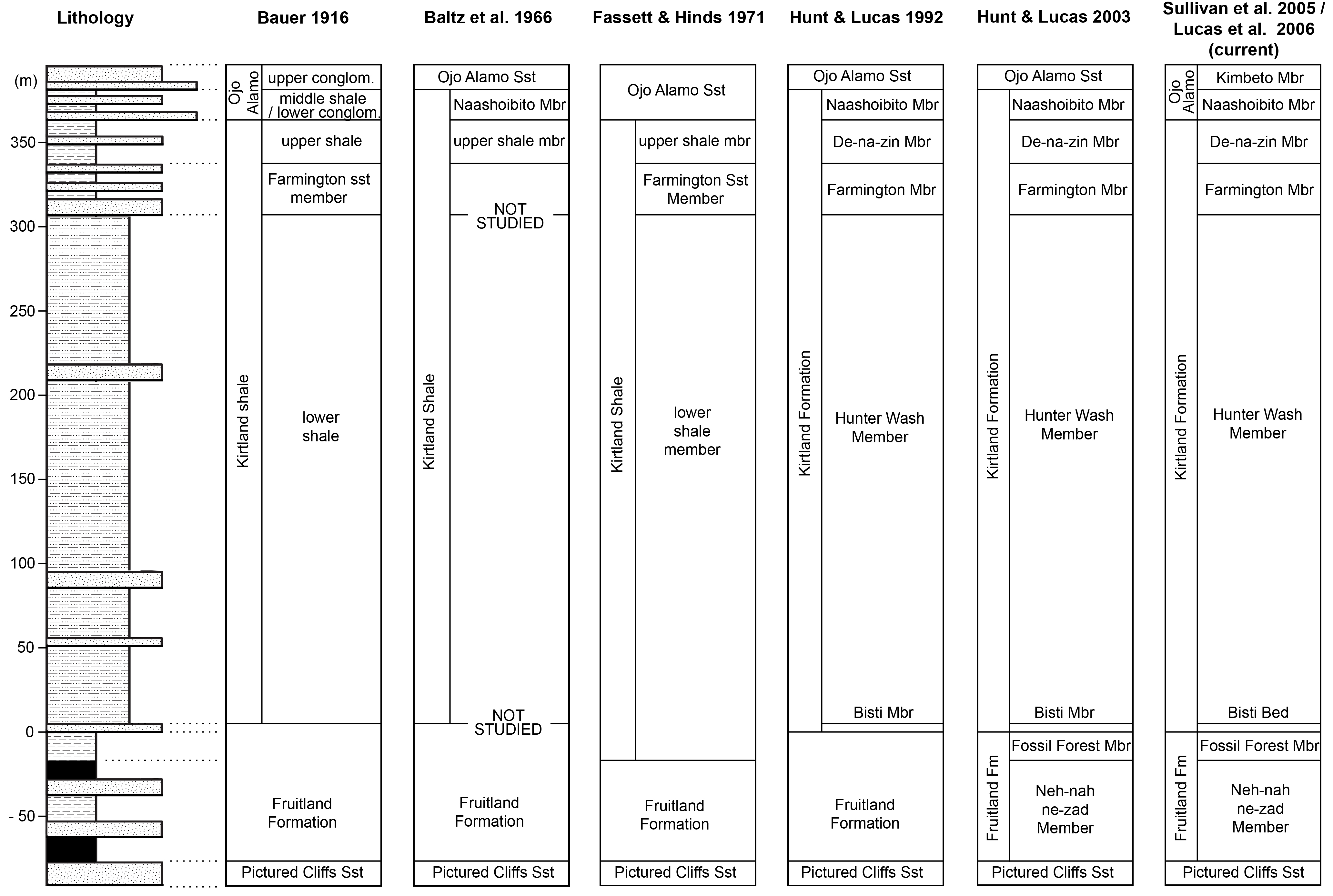

### Figure S4

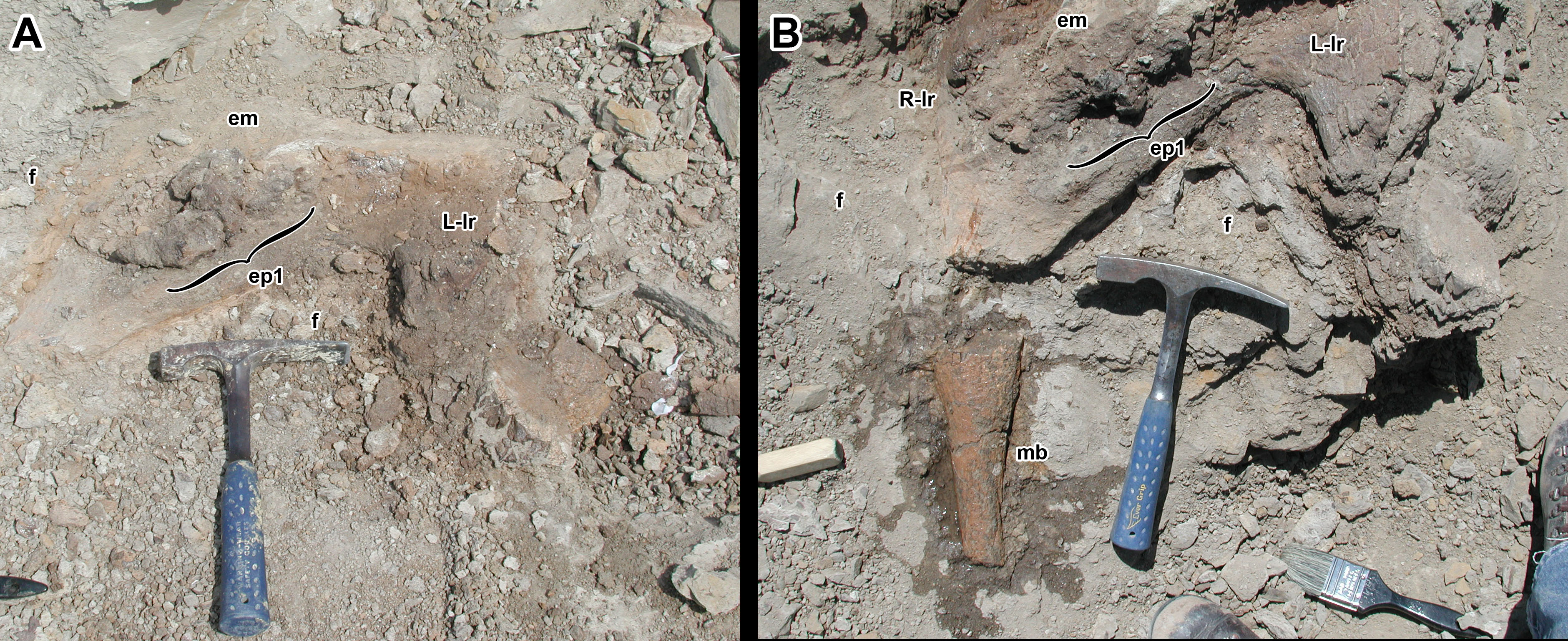

### Figure S5

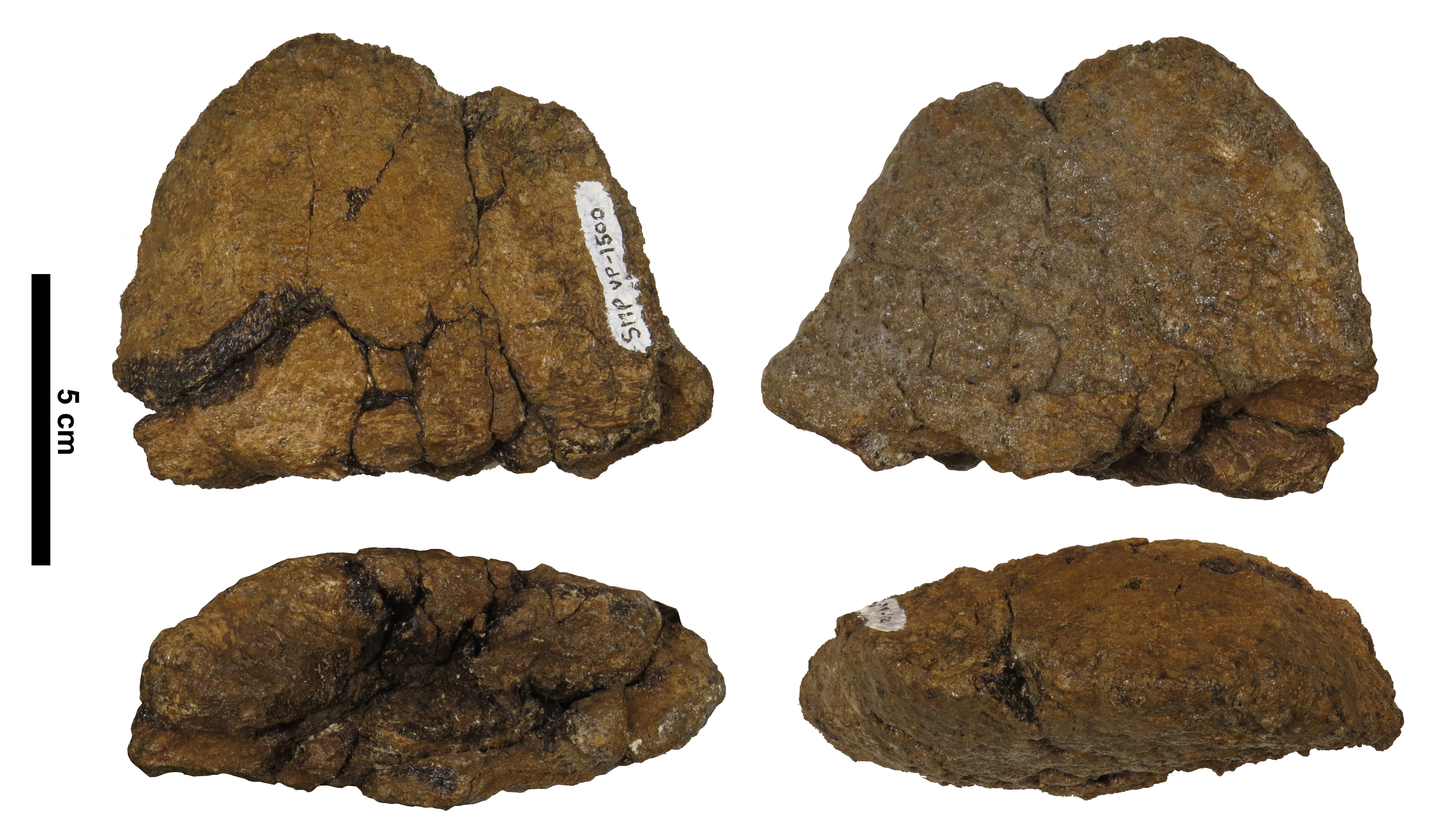

### Figure S6

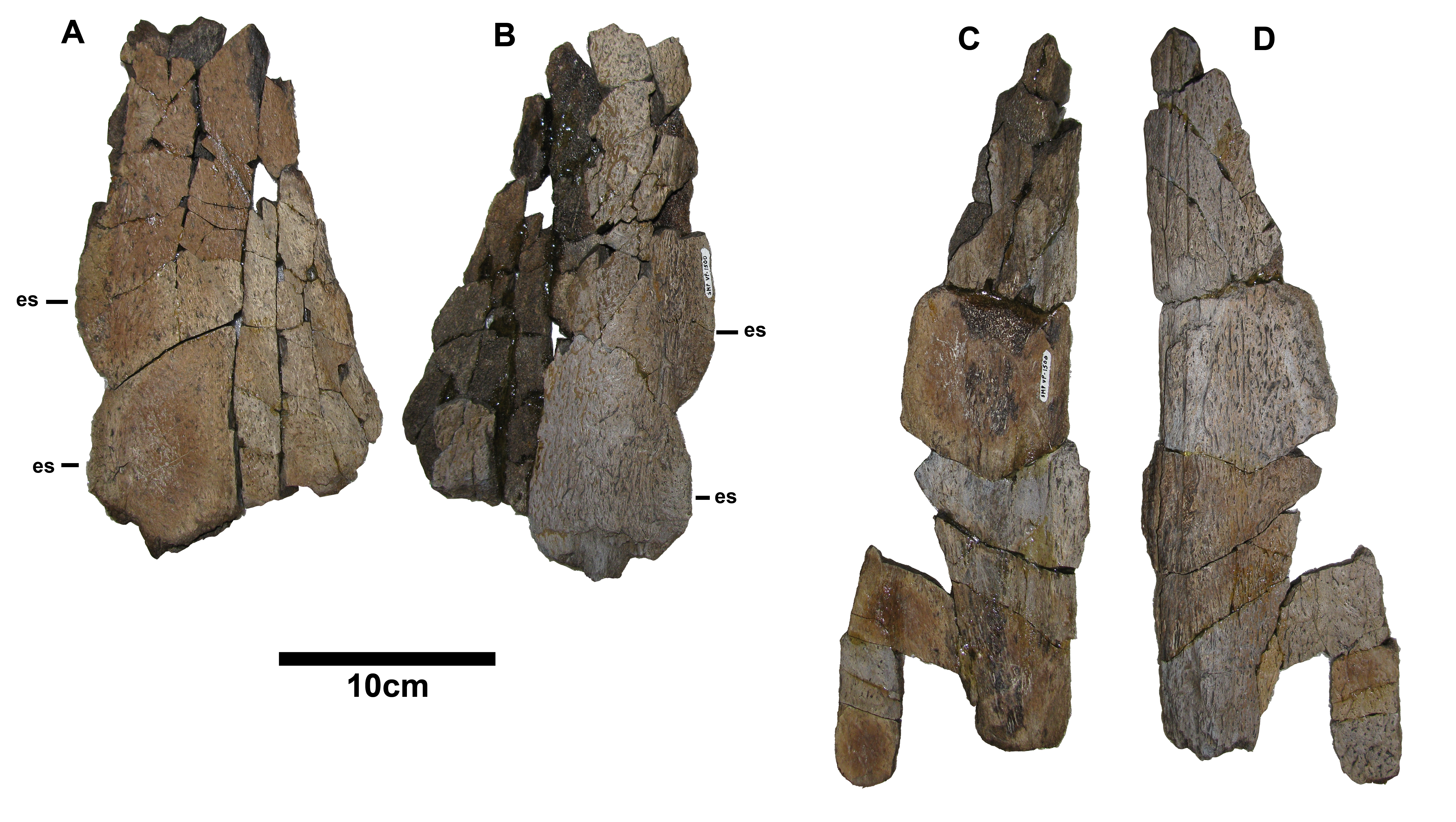

### Figure S7

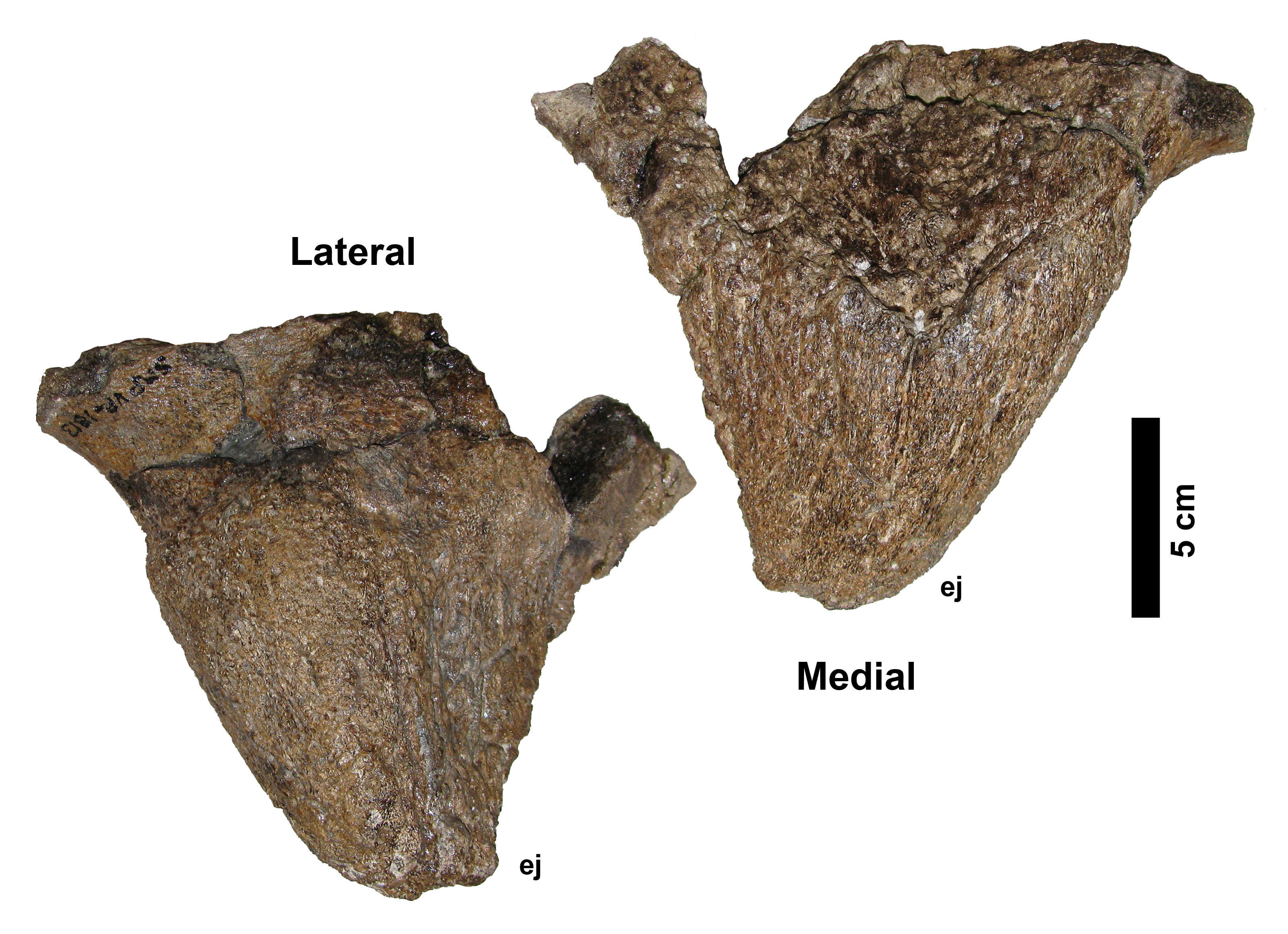

### Figure S8

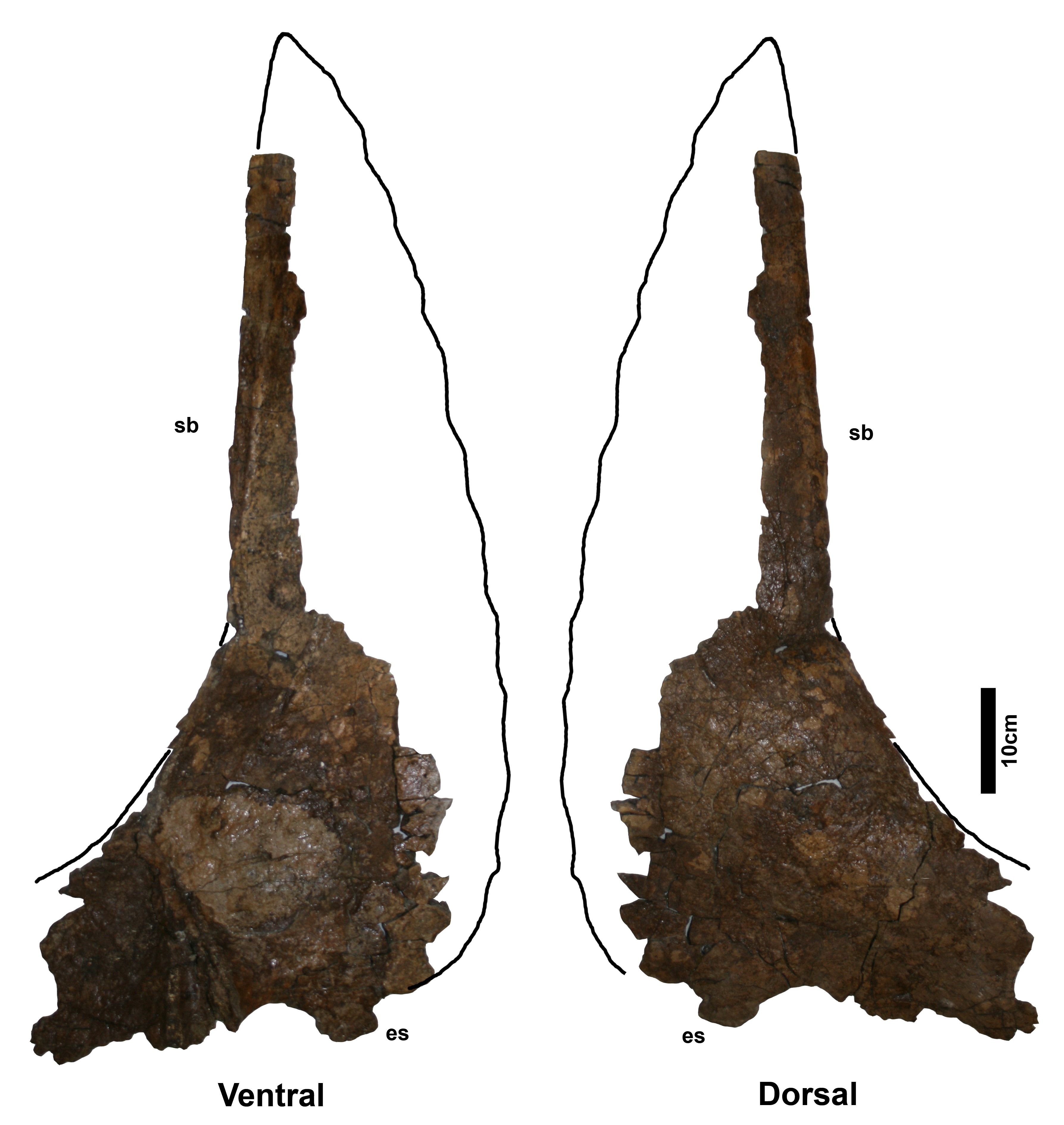

### Figure S9

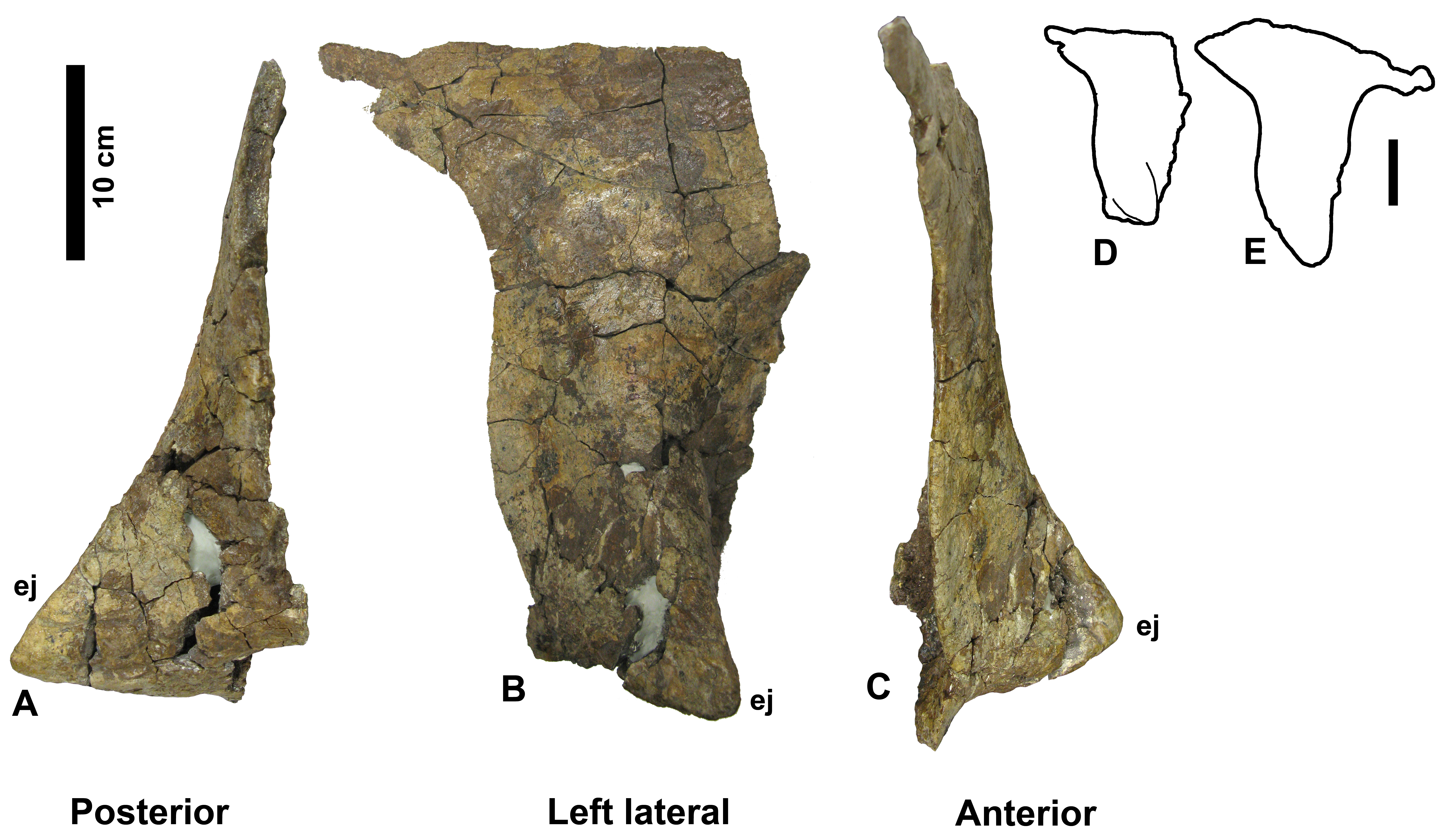

### Figure S10

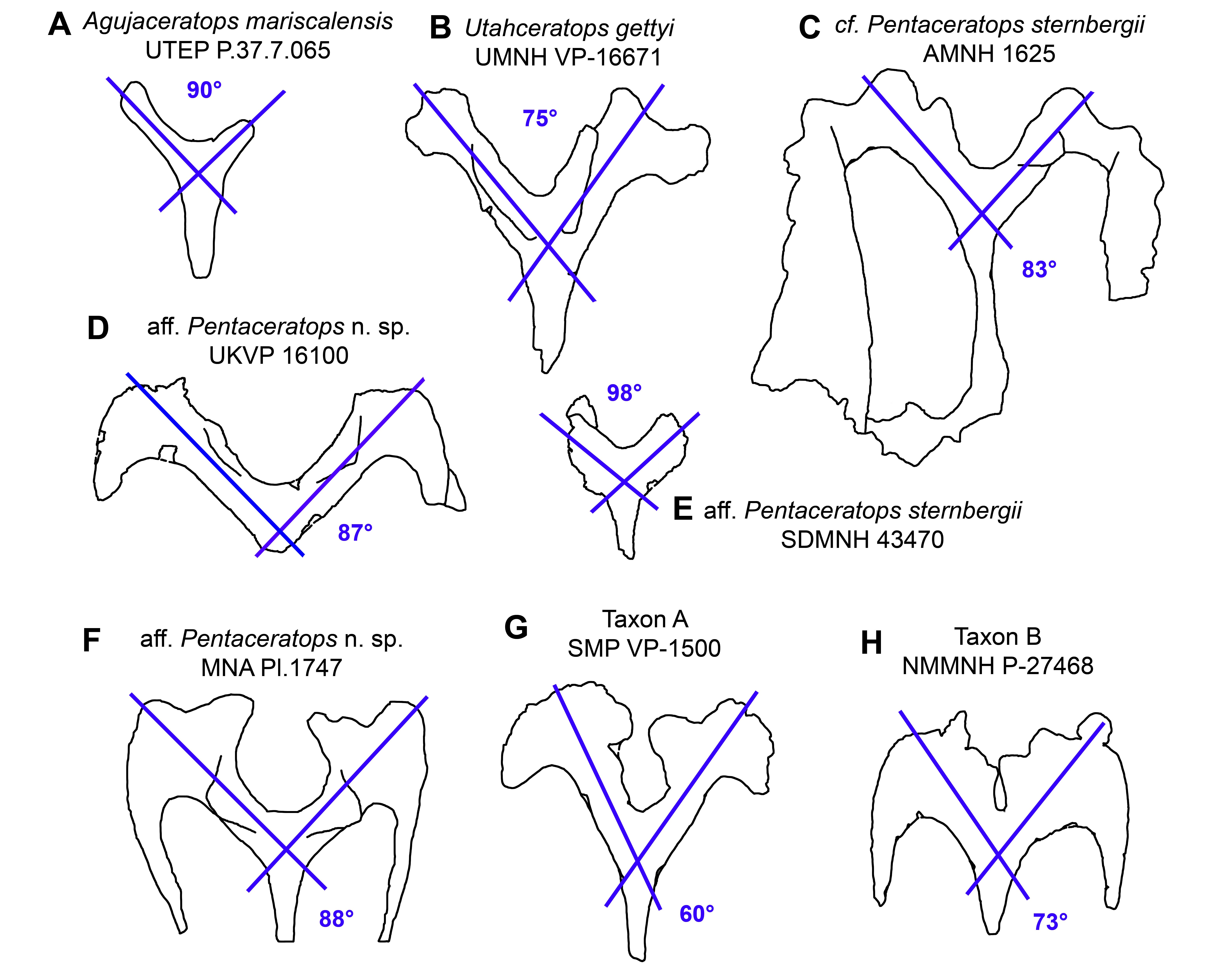

### Figure S11

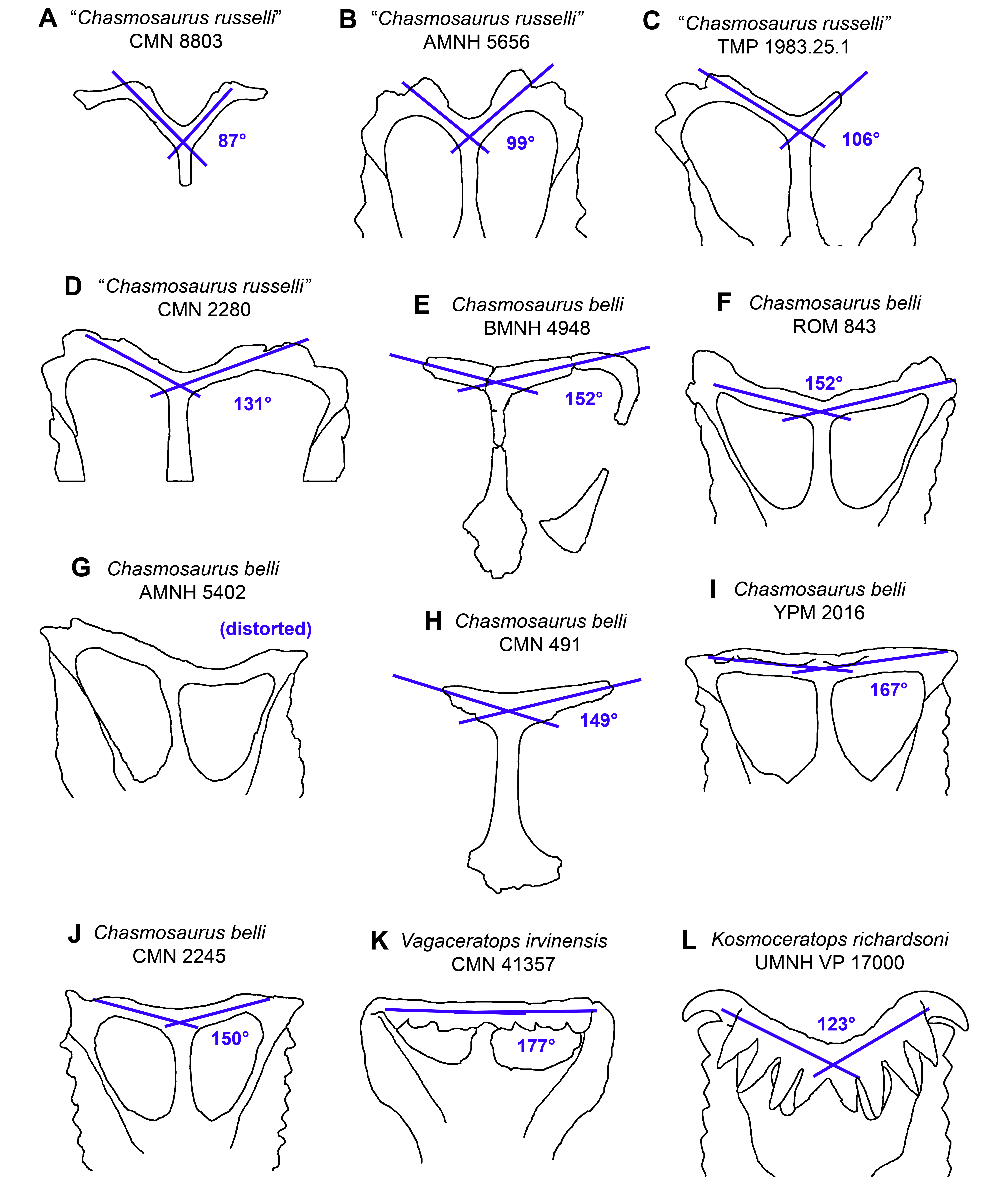

### Figure S12

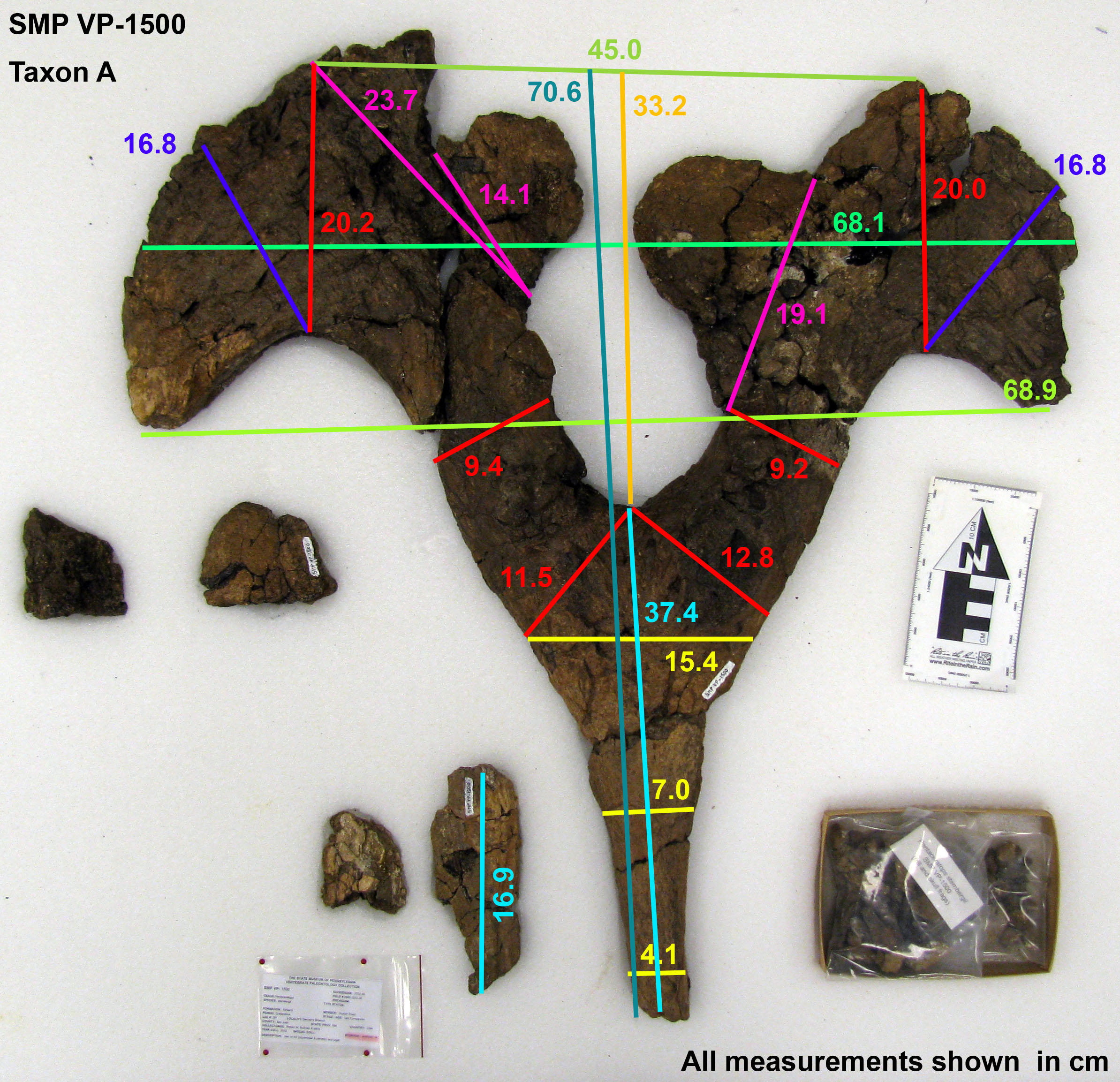

### Figure S13

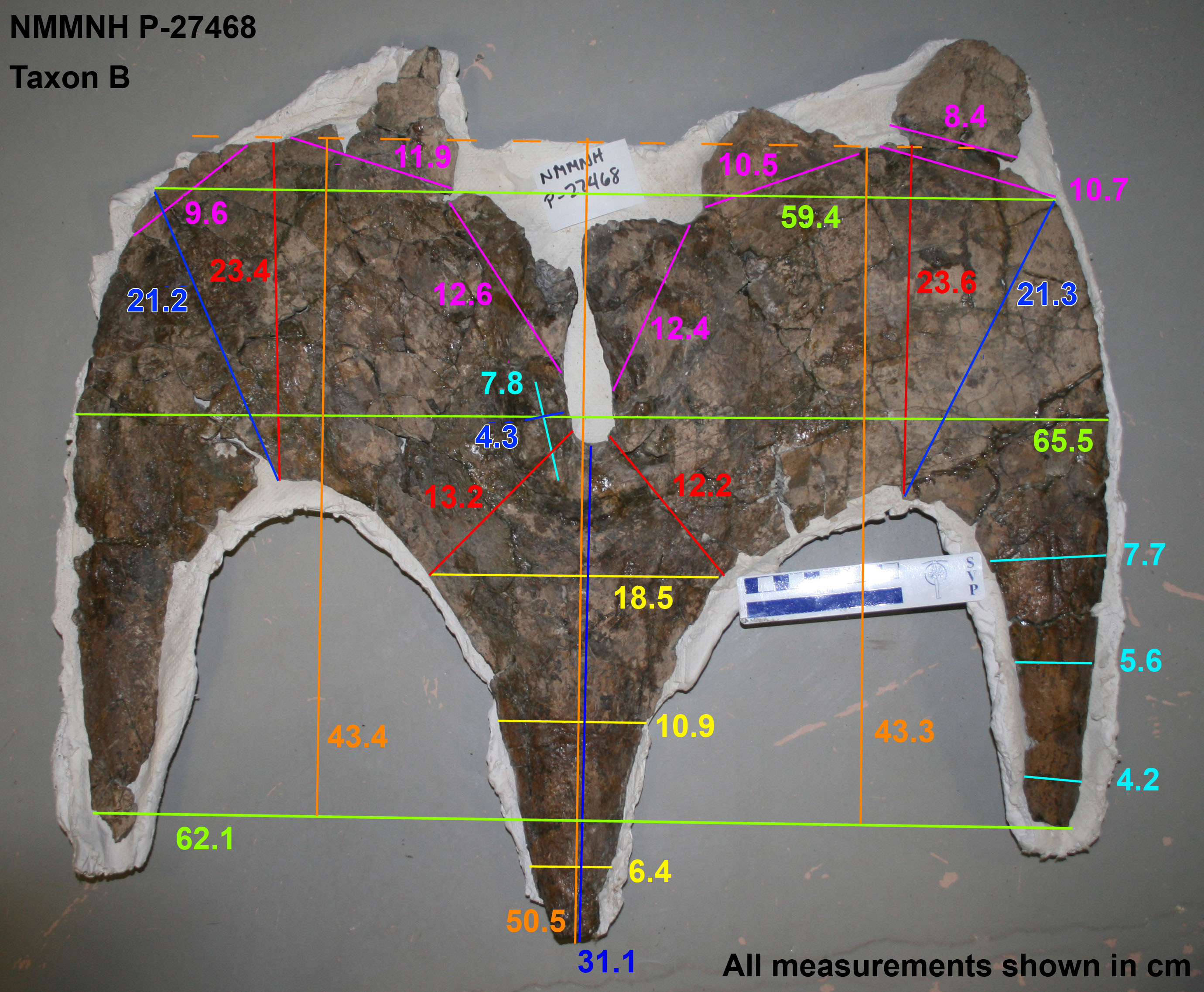

### Figure S14

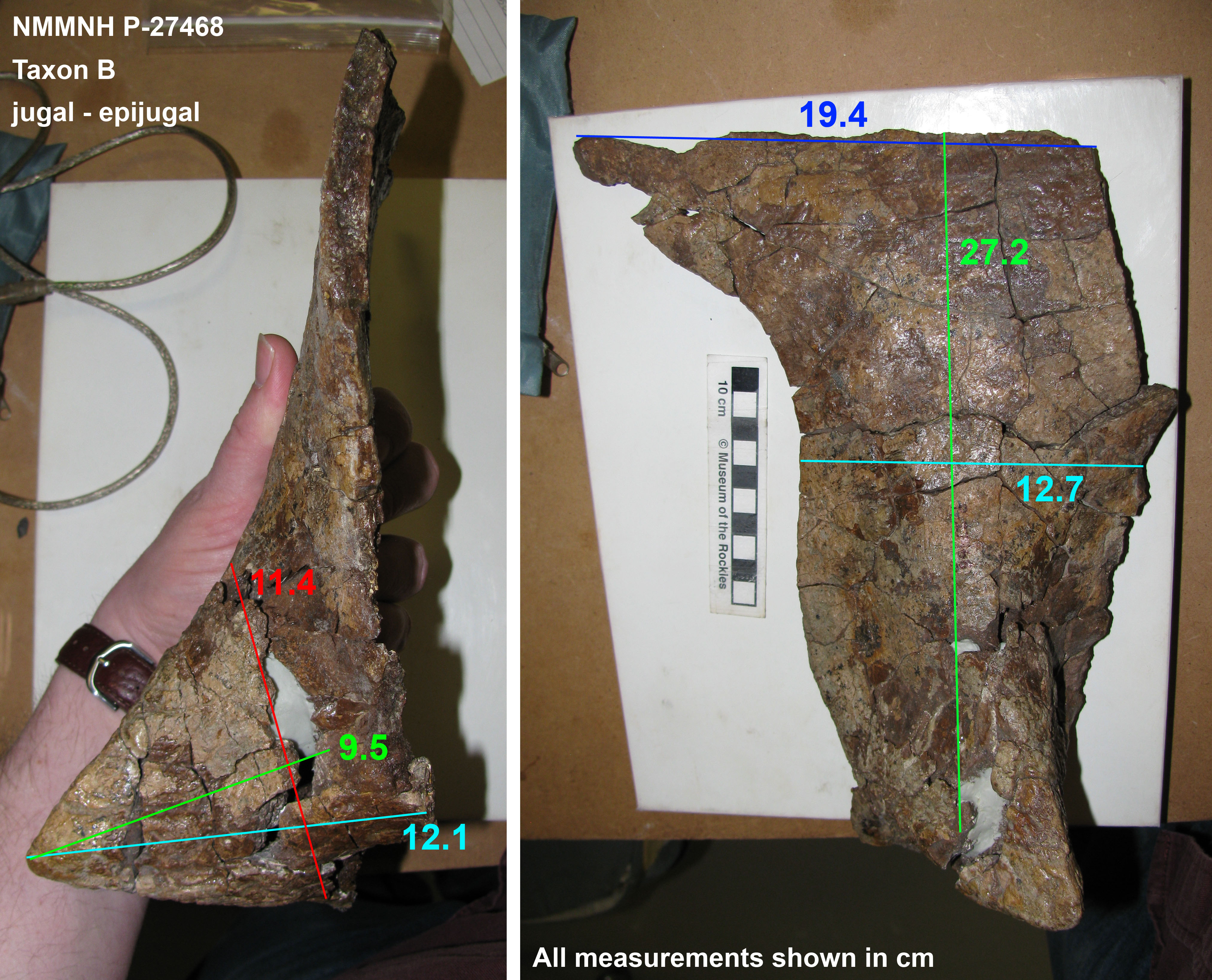

### Figure S15

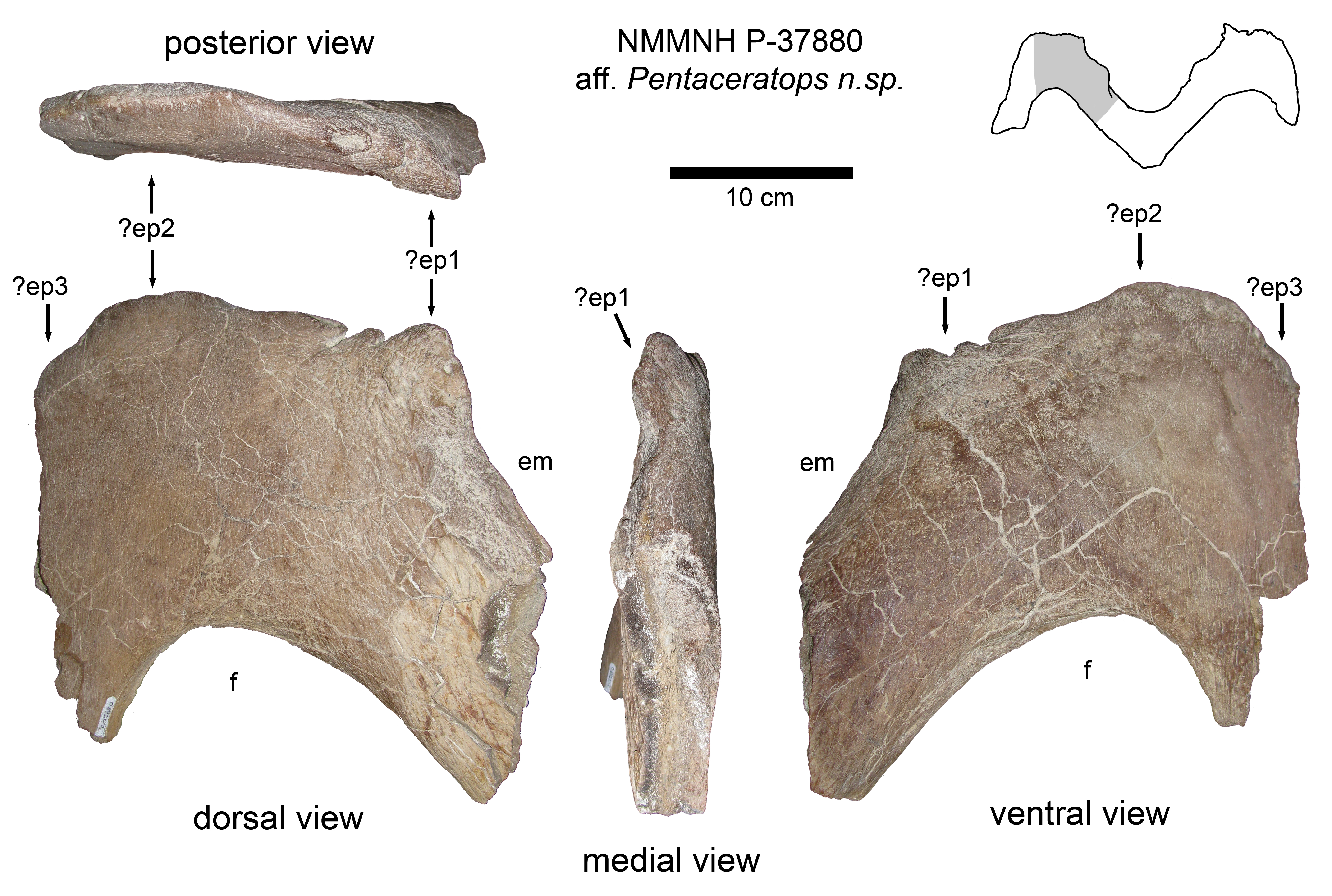

### Figure S16

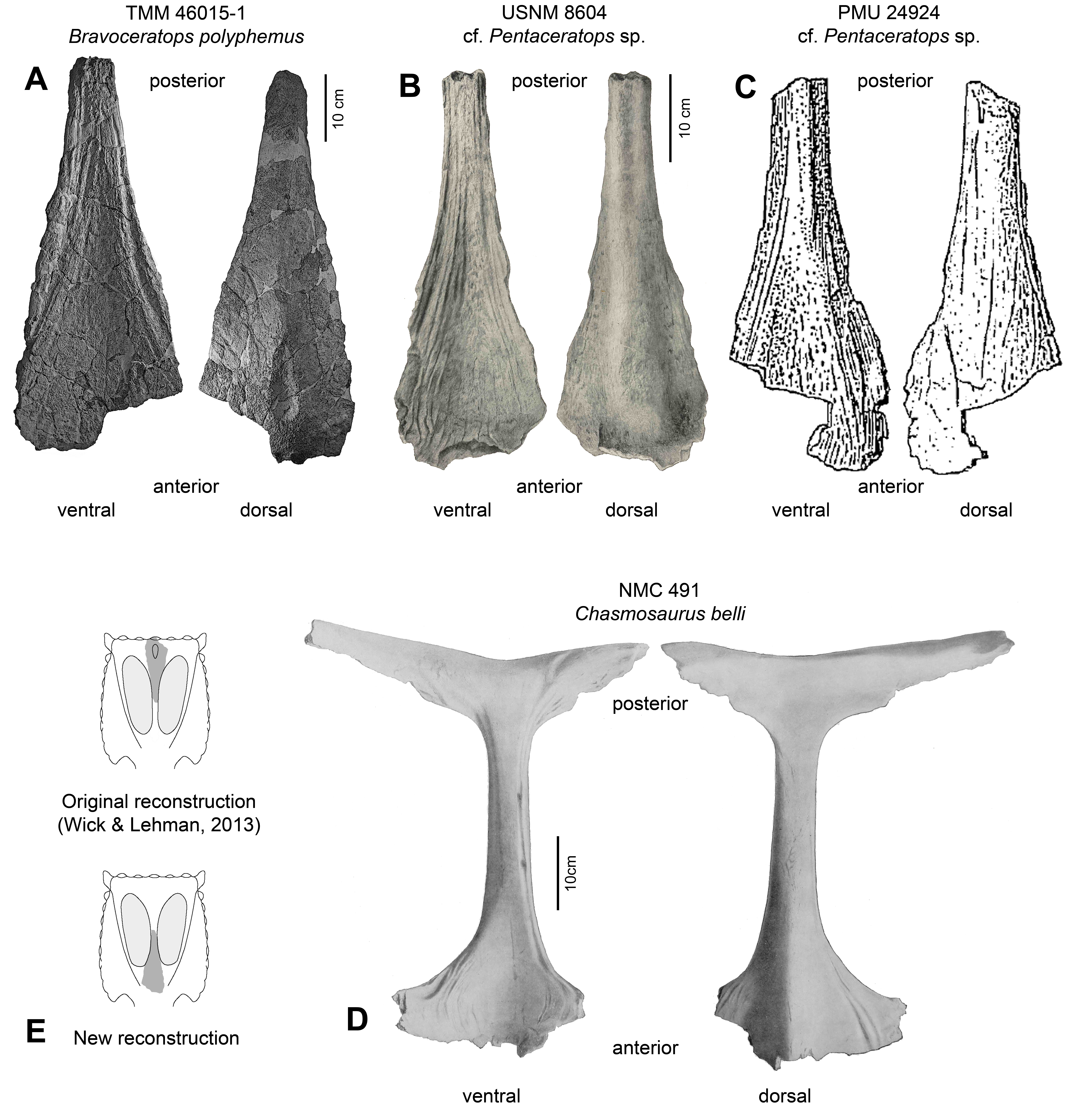

### Figure S17

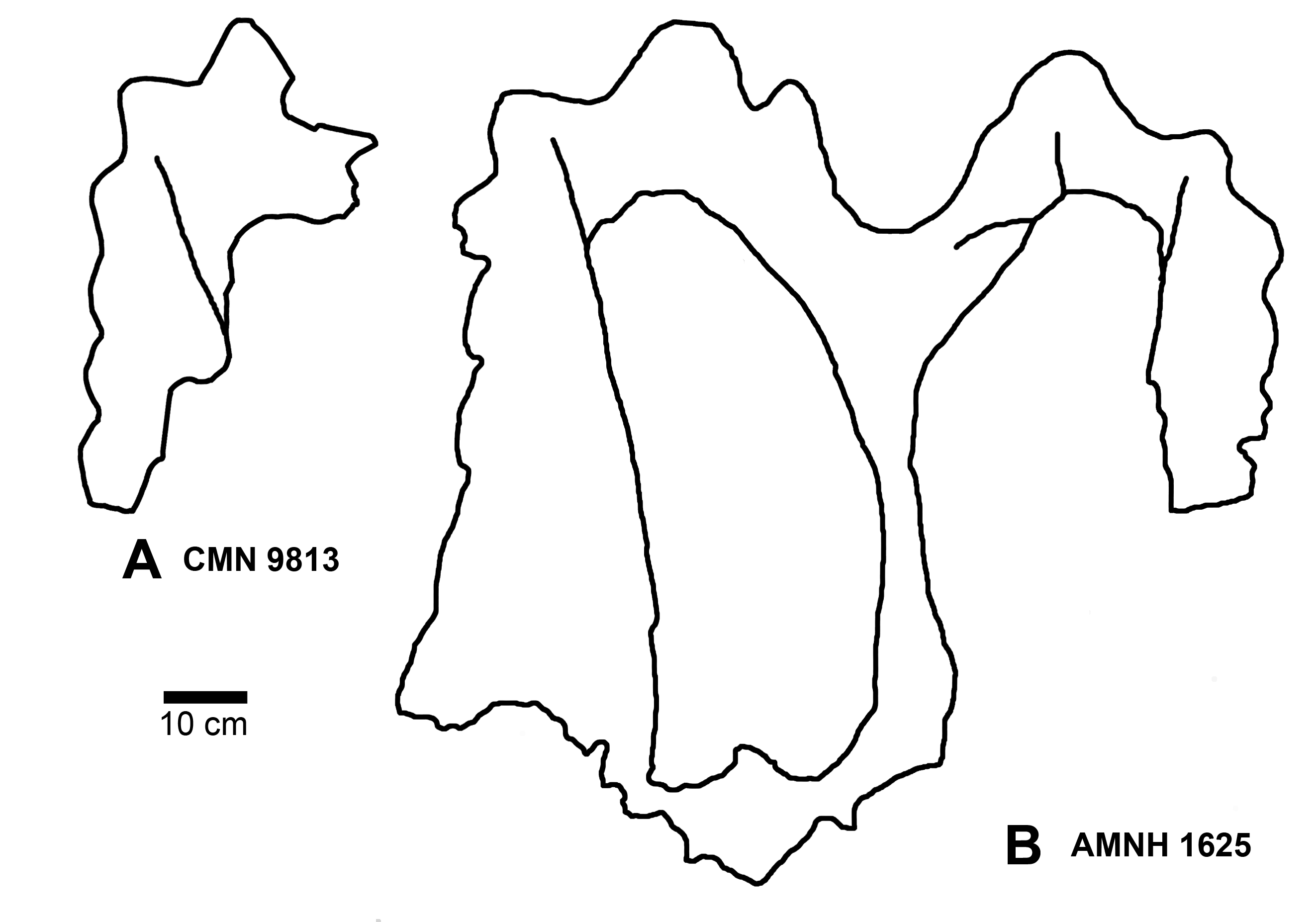

### Figure S18

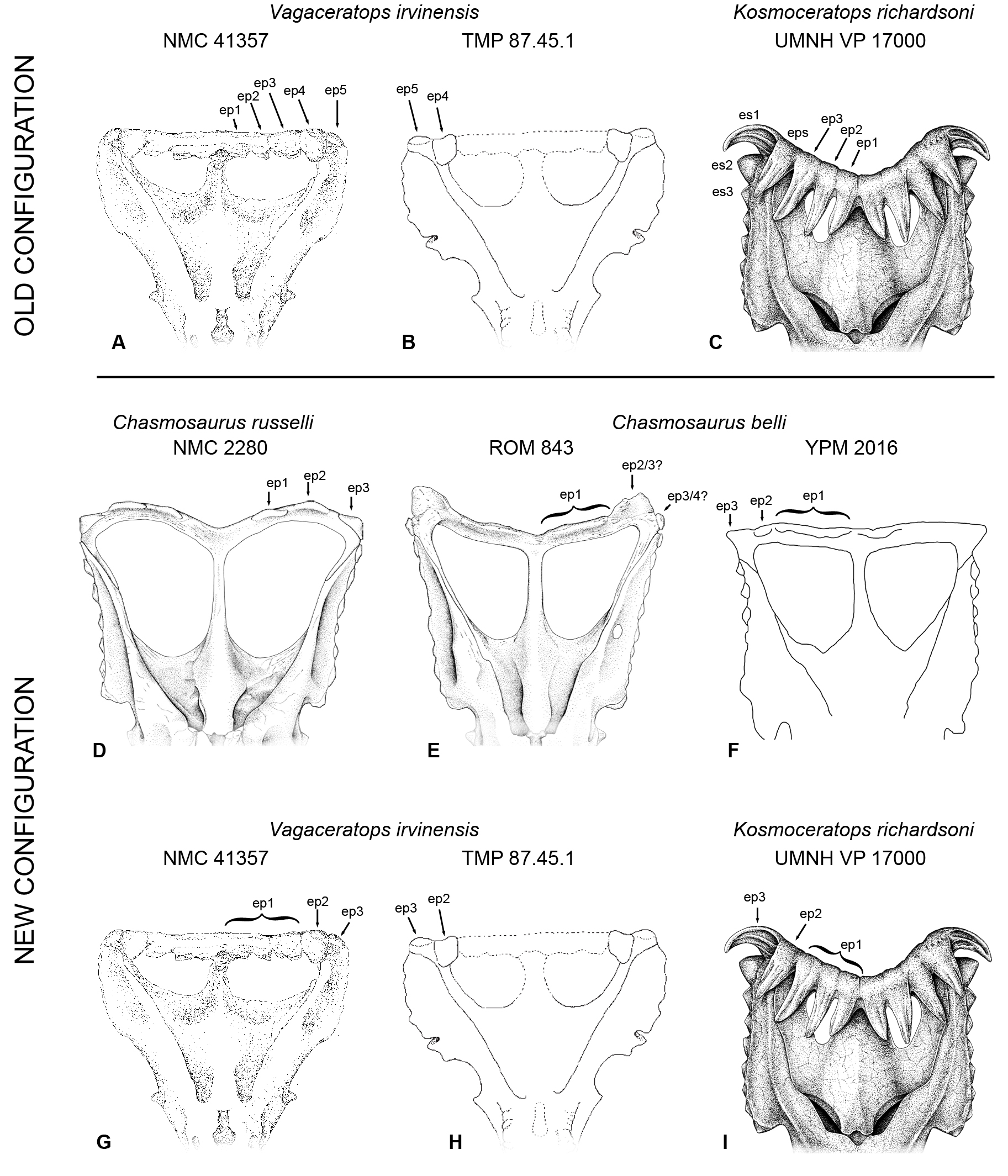

### Figure S19

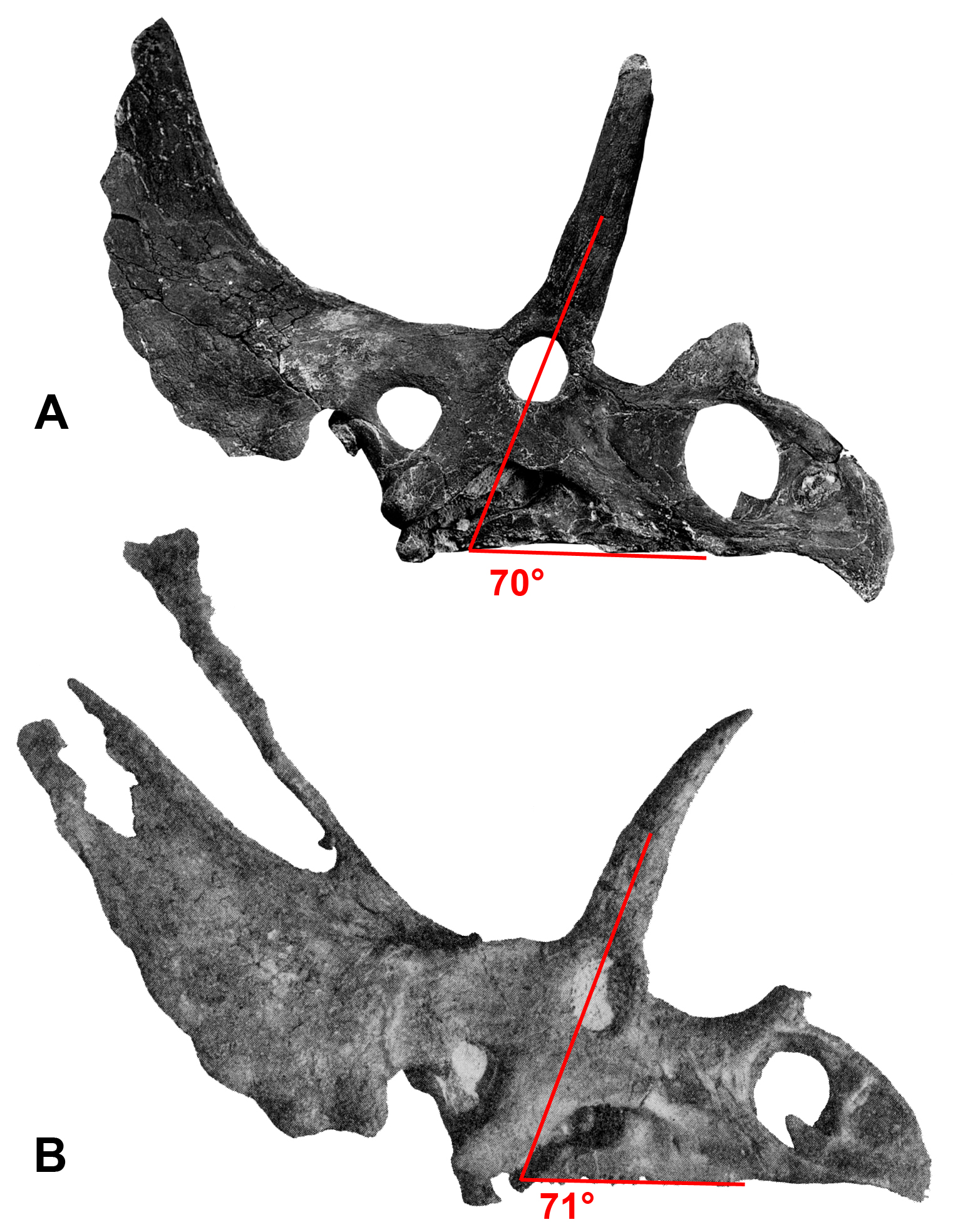
